# DNA Staples: An oligonucleotide library for data storage and computing

**DOI:** 10.64898/2026.05.11.724305

**Authors:** Kaya Selina Wernhart, Mathias Orlando, Fabian Schroeder, Ivan Barišić

## Abstract

DNA offers exceptional information density and stability, making it a promising medium for long-term data storage. However, the high cost of DNA synthesis and data retrieval remain key barriers to large-scale deployment. In this paper, we present DNA staples, a multipurpose library of short single-stranded DNA sequences that enables the encoding of arbitrary digital data by enzymatic assembly. Flexible encoding schemes allow the same presynthesized strand library to be used across applications, significantly reducing synthesis requirements while supporting diverse data representations. Using a restricted library also confers inherent error correction. In addition to storage, the library enables creation of computational DNA modules that perform highly parallel operations directly on stored data. This framework provides a cost-efficient approach to molecular data storage and supports integrated storage-computation at the DNA level.

## Introduction

DNA is a very promising medium for cold data storage as it can remain stable for millennia under suitable conditions [1, 2, 3]. Furthermore, it offers data densities that significantly exceed those of conventional storage [4, 5, 6]. In practice, storing data in DNA requires the physical construction of DNA strands encoding the desired information, achieved through different synthesis strategies. DNA synthesis currently relies on two main approaches: *base-by-base synthesis* or *block assembly*. In base-by-base synthesis methods, sequences are written nucleotide by nucleotide using chemical or enzymatic methods [7, 8, 9]. This approach maximizes flexibility in sequence design but accumulates errors during long-strand synthesis and remains costly, as each dataset requires *de novo* chemical synthesis of DNA strands [10, 11]. By contrast, block assembly enables the synthesis of kilobase long DNA molecules by assembling shorter, presynthesized DNA fragments from a predefined library. This method relies on biological DNA repair mechanisms to mediate the assembly of desired DNA fragments and is widely established as a standard strategy in gene synthesis [8, 12, 13]. Previous block assembly strategies in DNA data storage have relied on libraries of double-stranded DNA elements, which limit inter- and intramolecular interactions by exposing only short specific sticky ends [14]. Conversely, single-stranded DNA blocks offer greater flexibility and can reduce the number of distinct strands required in a library, making them an efficient choice for DNA data storage [15, 16]. However, single-stranded approaches are more sensitive to inter- and intramolecular interactions, particularly in highly repetitive sequence contexts such as those arising from limited libraries [17, 18]. Repetitive sequences are a known challenge in gene assembly and initial mitigation strategies have been explored, using double-stranded DNA building blocks [19, 20]. In this work, we introduce DNA stapling, a strategy of single stranded DNA block assembly. It is based on the assembly via partially overlapping hybridization (Figure 1b). DNA stapling relies on a single stranded DNA library that minimizes the number of individual blocks needed for synthesis of arbitrary data. This minimal library enables flexible design of encoding schemes and supports basic computational elements. Similar hybridizing libraries have been used previously, for various combinatorial problems such as the Hamiltonian path problem [21, 22, 23]. These suggest that our library can be extended to general computational approaches.

**Figure 1.**
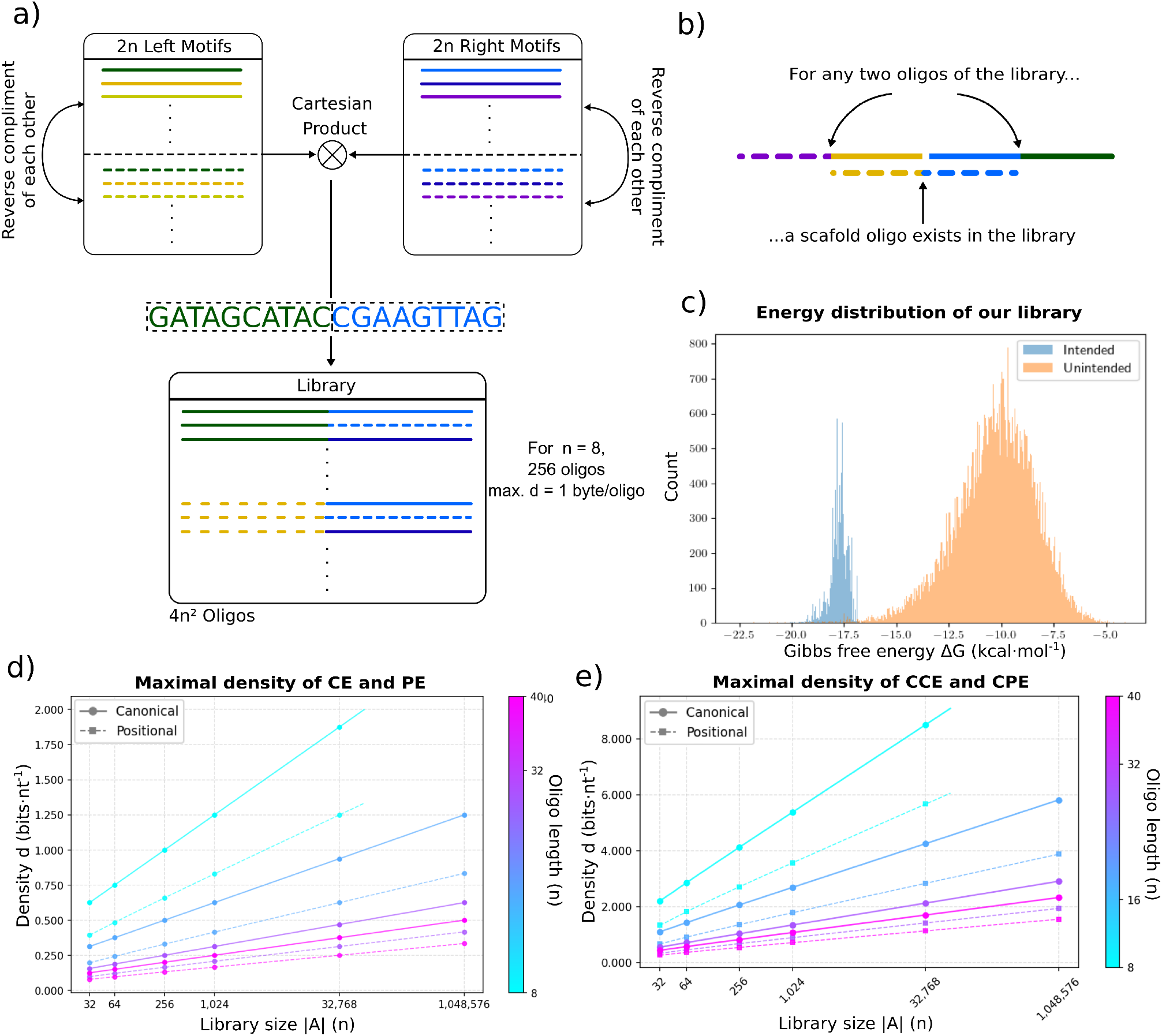
Construction and thermodynamic analysis of the DNA strand library. a) Strand libraries generated as the Cartesian product of motif domains, illustrating the relationship between domain set size and resulting library size. b) Schematic of scaffold-mediated hybridization between two corresponding oligonucleotides; each oligonucleotide pair is associated with a unique scaffold strand. c) Histogram of the highest-energy conformations for all strand pairs, comparing intended hybridization pairs with unintended interactions. d) Data densities for libraries of different oligo lengths and library sizes for the canonical encoding and positional encoding. The canonical encoding density is equivalent to the maximal density of a library. e) Data densities for libraries of different oligo lengths and library sizes for combinatorial encodings.

Our predefined DNA library addresses key limitations of current DNA data storage and molecular computation systems. Existing chemical synthesis based approaches scale linearly with data size because each dataset requires *de novo* oligonucleotide production, making large-scale storage costly. In contrast, our library was synthesized once and reused across arbitrary datasets, shifting the bottleneck from chemical synthesis to programmable assembly. The use of standard desalted oligonucleotides further reduced cost.

Reliable assembly requires predictable interaction dynamics within a partially complementary oligonucleotide set. Intended hybridizations were designed to be energetically similar and stronger than unintended interactions, minimizing off-target binding and enabling more controlled self-assembly. Oligonucleotides were designed with defined pairwise edit distances, a measure of sequence dissimilarity, which structured the interaction space and provided inherent error-correcting capability, enabling accurate recovery despite synthesis, assembly, and sequencing errors.

Data recoverability depends on encoding strategy, biochemical assembly dynamics and bioinformatic optimization. In our design, encoding was implemented through varying selections and assemblies of library elements into strands. We show that encoding choice strongly influenced both data density, recovery robustness, and assembly complexity, with clear differences observed across encoding schemes.

The library is further extended beyond data storage through strand displacement logic gates constructed from concatenated library elements. These gates implemented AND and OR functions, demonstrating that a single predefined library can support both information storage and molecular computation.

## Results and Discussion

### Library Design

Our DNA staple approach centers on a *programmable single-stranded DNA (ssDNA) library*, consisting of fixed-length ssDNA sequences called *oligonucleotides (oligos)*. The library architecture relies on defined hybridization domains at each end. Each oligo features a left binding domain at its 5’ end and a right binding domain at its 3’ end (Figure 1a). DNA’s antiparallel structure dictates domain-specific hybridization. Left domains hybridize only with left domains of other oligos, while right domains hybridize only with right domains. This domain exclusivity allows independent selection of left and right motifs for the library.

Any two library oligos assemble via ligation when a third oligo serves as scaffold. For every left motif *L*_*i*_ and right motif *R*_*j*_, the library contains an oligo with left domain reverse complement to *L*_*i*_ and right domain reverse complement to *R*_*j*_. Such libraries are generated from the Cartesian product of *l* left motifs and *r* right motifs. Each domain set includes its motifs plus their reverse complements, excluding self-complementary sequences. The inclusion of reverse complements in domains guarantees the existence of staples for all oligo pairs. The design generates libraries of size *rl* with data

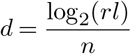

bits per nucleotide for oligo length *n* (Figure 1d). The choice of *n* therefore reflects a trade-off between data density, edit distance and hybridization specificity.

Scaffolds hybridize to two adjacent oligos and position them for enzymatic ligation into continuous double-stranded DNA (dsDNA). To achieve assembly in arbitrary order while minimizing total library size, each library oligo has to serve as both a data-encoding element and structural scaffold. The dual role demands uniform library architecture: all oligos share identical lengths and domain lengths. The library thus supports universal hybridization between arbitrary oligo pairs to assemble long dsDNA strands. A schematic of the hybridization is found in Figure 1b. Our library design achieves full programmability and maximal data density, as every possible ordering of library oligonucleotides can encode unique data using only library elements. This strategy extends previous work on scaffold-assisted DNA assembly, by combining both data-encoding and structural functions into each oligo [16]. Furthermore, the library structure lacks explicit encoding-dependent structural components. It therefore provides greater flexibility and allows more variation in encoding strategies compared to other block-assembly libraries [14, 24].

For implementation, the library requires high hybridization specificity. Off-target binding can block intended strand assembly. We developed a two-step screening process to select optimal motifs.

The first step restricts candidate DNA sequences to a smaller set with uniform thermodynamic properties. Left and right motifs therefore have identical lengths and GC-content. This design ensures consistent hybridization energies across all pairs. Screening further excludes palindromic sequences that cause hairpins and self-reverse-complement motifs that block domains.

From these motif pools, the second step constructs test libraries by randomly selecting motifs such that the set of motifs has at least edit distance *d*. Using NUPACK, we compute binding energies between oligo pairs of a respective test library [25]. Intended pairs are those that have at least one motif that is reverse complementary to each other. These pairs should exhibit higher binding energies than non-intended pairs. We use the largest difference between the minimal intended and maximal unintended binding energies as a library selection criterion (Figure 1c).

The library used in this work comprises of 256 oligonucleotides each 20 nucleotides (nt), consisting of 16 unique left and 16 unique right motifs of length 10 nt. Left and right binding domains have 50% GC-content, with edit distance 3 between all of them. This enables 1 byte of data storage per oligo as log_2_(256) = 8 bits, at a maximum data density of 0.4 bits per nt. The full sequence library is listed in Supplementary Table S5.

### Encoding Strategies

Because most DNA data storage approaches are based on nucleotide by nucleotide synthesis, they encode information into sequences over the canonical four letter alphabet (A, C, G, T). They maximize data density with respect to sequence-level error correction and biochemical constraints, bound by the theoretical maximum of two bits per nucleotide [26]. Recent work has also extended this approach by exploring composite alphabets that mix nucleotides to form a larger effective symbol set, thereby increasing logical density per synthesis cycle to above 2 [27, 28]. Preuss et al. formalize shortmer-based combinatorial alphabets whose symbols consist of subsets of distinguishable short DNA oligomers, leading to a significant increase in logical density. Similarly, composite DNA studies demonstrate that using expanded alphabets can achieve higher bits per symbol while still employing error-correcting codes tailored to these extended alphabets [29].

In comparison, block assembly based approaches design oligonucleotide libraries to implement a predefined encoding strategy [16, 14, 15]. Here, we invert this construction by first defining the block library and subsequently developing encoding schemes that operate within it. Our library allows for various encodings and assembly strategies to encode data. In this respect, our schemes generalize the traditional nucleotide-based encoding beyond a quaternary alphabet to an n-long library alphabet. In *de novo* synthesized DNA data storage, each encoding scheme fixes the data density of the stored dataset [5, 4, 30, 3]. In contrast, the approach here constructs a combinatorial library in which the maximal data density depends on the number of oligonucleotides in the library (Figure 1d,e). As the number of library elements grows, a larger fraction of the possible sequence space becomes occupied, which generally reduces the minimum edit distance between sequences in the library, decreasing inherent error resilience. This trade-off between library size and error tolerance is well known in classical error-correcting code theory, where the minimum distance between codewords determines the number of errors that can be reliably detected or corrected [31, 32, 33]. Similar considerations apply to all error correcting DNA data storage systems, where valid sequences must be distributed in sequence space to ensure reliable decoding under realistic synthesis and sequencing error rates [34, 35, 5, 36].

Further, assembly constraints, if used in an encoding, directly impact encoding choices, giving rise to error correcting codes tailored to the assembly errors. We define multiple *encoding schemes* that enable flexible data encoding. These schemes redefine data distribution across oligos and define DNA strand assembly strategies to optimize correct assembly and yield. Each approach involves trade-offs between storage density, assembly yield, and error correction capability. We examine four specific implementations in detail.

#### Canonical encoding (CE)

The canonical encoding, depicted in Figure 2a, maximizes data density for a single assembled DNA strand. The encoding process operates as follows: input data represented as bits are split into smaller bit sequences of length ≤ log_2_(|*A*|), where |*A*| is the library size. Each bit sequence maps to a unique oligo from the library, yielding an ordered list of oligos that define the data encoding DNA strands of the final assembled strand. The complementary strands are uniquely determined by the library construction and serves as scaffold between adjacent data-encoding oligos. Ligation of this ordered oligo sequence produces the final continuous encoded strand.

**Figure 2.**
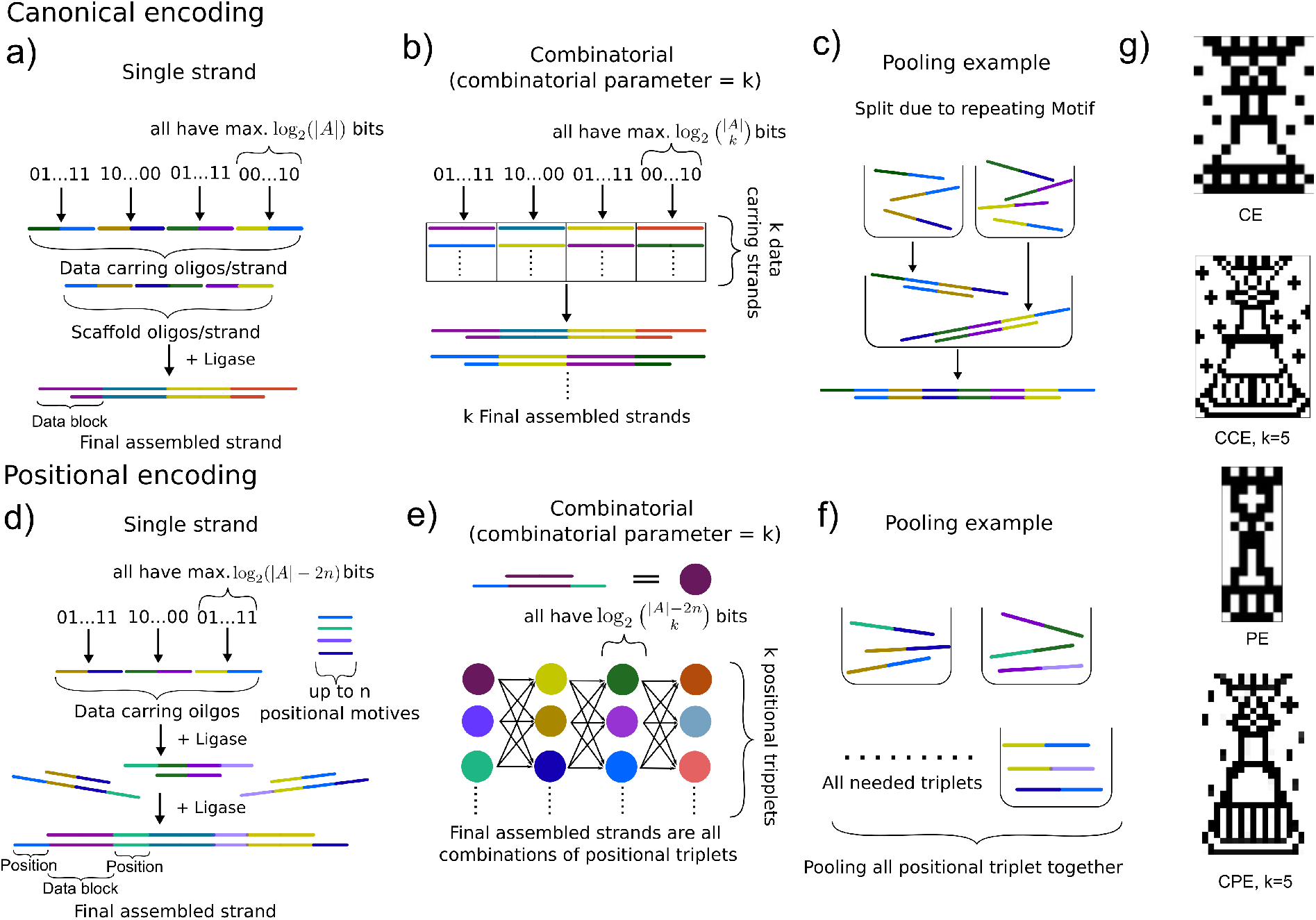
Canonical and positional encoding strategies for data-encoding DNA strands. In all panels, strands and their complementary strands are shown in identical colors. a) Construction of a single data-encoding strand using the canonical encoding scheme, illustrating motif hybridization and highlighting the final data-encoding block within the assembled strand. b) Combinatorial selection of data-encoding oligonucleotides in the canonical encoding scheme and their assembly into complete strands. c) Schematic depiction of the ligation process in canonical encoding, showing the requirement for separated oligonucleotide pools prior to final assembly due to repeating motifs. d) Construction of a single data-encoding strand using positional encoding, illustrating triplet formation and the distribution of structural and data domains. e) Combinatorial assembly of positional encodings, enabling strand-blind assembly and increased data density without additional synthesis cost. f) Schematic depiction of the two-step assembly process required for positional-encoded strands. g) Images encoded using the canonical and positional encoding schemes.

The assembly protocol, the stepwise assembly of oligos into final DNA strands, depends on the specific encoded data due to inevitable motif repeats within the ordered oligo sequence. An example of an assembly protocol is shown in Figure 2c. Repeated left or right motifs cause ambiguous intended bindings because any oligo with a matching motif can bind regardless of its intended position in the sequence. For each data strand there exist multiple assembly protocols and selecting one can impact assembly errors and final yield. Each ligation step comprises all parallel ligation pools defined by the assembly scheme and performed in parallel at that step. By design, each primary pool contains at least three oligonucleotides.

#### Positional encoding (PE)

The positional encoding minimizes ligation steps to only two and renders the assembly protocol data independent. However, this comes at the cost of reduced data density. This scheme defines *positional triplets*, groups of three oligos ligated separately. These are then combined to the final strand in the second step (Figure 2d).

To encode data, PE uses two unique positional motifs per position, one from each domain. Adjacent positions use reverse complements as positional motifs to ensure linking. These motifs remain unique within each pool, ensuring unambiguous hybridization in the final ligation step. The second ligation joins the positional triplets into the final strand in one reaction.

Each triplet includes one *data-encoding oligo* selected from a restricted library excluding all oligos that include positional motifs or their reverse complements, at any given position. These data-encoding oligos determine two *scaffold motifs* that, together with the positional motifs, determine the last two oligos completing the triplet. These restrictions limit data-encoding oligo choices and reduce data density. The encoding process splits input bits into smaller bit sequences of length ≤log_2_ (|*A*|*− r− l*), assigns them to triplets, and yields an assembly protocol determined entirely by its construction.

This encoding limits the maximum number of positions per codeword to 2 *·* min(*l, r*) *−*1. For longer strands, extension strategies construct a second strand that assembles to the last positional motif and adds an additional ligation step using position-shifted triplets, enabling continuous strand formation.

#### Combinatorial encoding

DNA data storage usually stores many copies of the same strand. If multiple variants of the same strand can be created at no additional cost, this can be exploited to extend the alphabet and thereby increase the data density. Combinatorial encodings exploit this by using *k* different data-encoding oligos distributed across different strands. The specific combination of these *k* oligos at each position encodes additional bits beyond single oligo selection. This is based on previous work of Preuss at al. [27].

This approach works for both encodings but benefits positional encoding most. Practically, this approach adds no cost only when different oligo combinations can form naturally during assembly. For PE this is possible. Positional motifs uniquely determine each position, ensuring any final strand assembly remains compatible regardless of specific data-encoding oligo combinations used. The resulting data densities per oligo are:

- Combinatorial positional encoding (CPE): 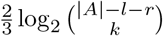 bits
- Combinatorial canonical encoding (CCE): 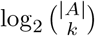 bits

The increase in density can for *k* = 5 is depicted in Figure 1e. Figure 2b and 2e show schematic representations of these combinatorial encodings.

### End-to-end data storage

To verify experimentally our library we encoded four black and white chess queen images (Figure 2g) using our 256-oligo library across the four encoding schemes with parameters from Table 1. For all encoding schemes, 51 oligos make up one dsDNA strand. All oligos were only salt purified to demonstrate low cost production is sufficient for this application.

**Table 1.** Encoding parameters for Queen images at four resolutions. All strands are 520 bp long (canonical) or 1560 bp long (positional triplets).

| Image | Bits | Strand length (bp) | Encoding | Combinatorial parameter $k$ |
| --- | --- | --- | --- | --- |
| Queen (lowest res.) | 119 | 520 | Positional | 1 |
| Queen (low res.) | 208 | 520 | Canonical | 1 |
| Queen (high res.) | 544 | 520 | Positional | 5 |
| Queen (highest res.) | 832 | 520 | Canonical | 5 |

Supplementary Algorithm S1,S2 provides pseudocode for these encoding calculations. Assembly protocols tailored to each encoding scheme enabled the construction of the four queen images. Supplementary Materials S7 detail all assembly protocols.

#### Purification and assembly verification

We tested all encodings under three conditions: no purification, one purification before final assembly and two purifications, one before final assembly and one to excise the final product. Gel excision purifies target strands from byproducts to analyze effects of unligated strands on the assembly. Assembly protocols ensured target strands reached 70 bp minimum length at purification, matching gel extraction kit specifications. Assemblies performed without purification yielded products with inconsistent strand lengths across all encoding schemes. Incomplete ligation created incomplete fragments acting as unintended scaffolds in subsequent ligation steps, generating diverse byproducts of various lengths. These errors propagate through every ligation step. A single purification step ensured assembly products that matched target length for all encodings except CPE. The second purification step ensures that the final assembled product is of target product length. Prior to the first purification step, assemblies across all encoding schemes produced target bands at the expected sizes for all pools. However, some pools hybridize inconsistently due to oligos in the library either failing to hybridize, blocking assembly through unintended interactions, or making larger unexpected concatemers, producing faint or smeared bands. Faint bands, thus, hint at defective library oligos.

While the library design can mitigate hybridization-driven byproduct formation, synthesis-derived errors remain a fundamental limitation. Oligonucleotide production itself is inherently error-prone [37, 38]. During synthesis, the coupling efficiency per nucleotide is typically 98.5–99.5% [39], corresponding in the best case to approximately 91% full-length oligonucleotides in the library. Truncation reduces domain specificity and ligation efficiency, thereby promoting incomplete or off-target assemblies. Similar effects have been reported in genome assembly, multiplex PCR, and hybridization chain reaction, where they are commonly mitigated using costly size-selection purification methods such as PAGE or HPLC [39, 40, 41]. By leveraging error-correcting capabilities inherent to biological systems, bioreactor-based production offers a potential route to reduce synthesis-induced errors during large-scale oligonucleotide manufacturing, thereby lowering overall costs [42].

The assembly products were blunted using ssDNA caps with the reverse complement motifs of the end motifs and sequenced via long-read sequencing. We observed that the size distribution of sequenced strands for each encoding scheme correlated with number of purification steps, where no purification or one purification provided ample material while two purifications yielded sparser sequencing coverage. Notably, CCE showed substantial material loss after the first purification, as strands assemble independently, requiring significantly more pools and therefore more purifications for assembly. These cumulatively lose material.

Agarose gels had confirmed target strand lengths. However, sequencing revealed incomplete nick ligation due to limited ligase efficiency. Throughout this work we used T4 DNA ligase, the most commonly used ligase in molecular biology. Patapov et al. reported ligation efficiencies for T4 ligase of approximately 80% for single nicks after 1 h at 25°C [43]. Each encoded image contained 49 nicks making complete ligation highly unlikely. These results indicate that enzymatic ligation efficiency represents a fundamental constraint for large multi-oligo assemblies, effectively introducing an additional biochemical source of errors in DNA storage systems.

Figure 3 shows the strand length distribution of sequenced and base-called DNA strands. Assuming that incomplete nick ligation occurs with equal probability at each position, the observed distribution corresponds to an average ligation efficiency of 55.1%. However, additional factors, such as distinct ligation steps, may influence this distribution and alter the probability of ligation at specific positions. These results highlight that the assembly process is imperfect and itself constitutes a source of error.

**Figure 3.**
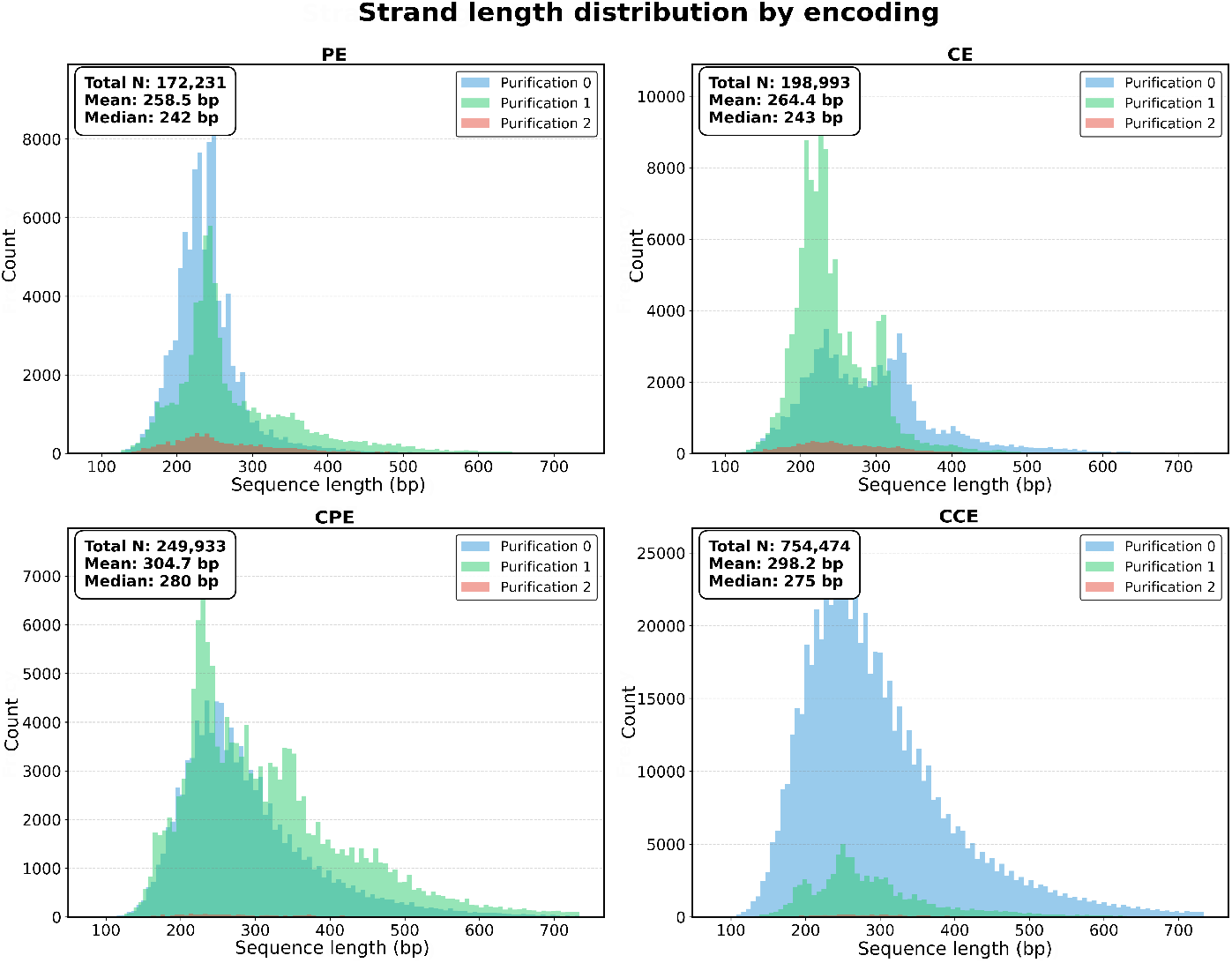
Distribution of recovered DNA strand lengths from nanopore sequencing across different encoding schemes and purification conditions.

Enzymatic ligation efficiency imposes a fundamental scaling limit on multi-fragment DNA assembly, resulting in exponential yield decay with increasing ligation steps. This limitation is particularly severe for repetitive or highly modular sequence architectures, where repeated concatenation of similar building blocks increases the number of required ligation steps and amplifies the probability of incomplete assembly. Consequently, ligation dependent systems such as synthetic genome construction are constrained to shallow assembly architectures, i.e., assemblies involving only a small number of sequential ligation steps, reducing the fraction of fully assembled strands.

#### Data recovery

Despite low ligation efficiency and error propagation, the data could be recovered successfully. Recovery begins by aligning the library oligonucleotides to the sequenced strands using the BLAST algorithm, retrieving chains of oligonucleotides that correspond to the assembled strands [44]. Two parameters control this reconstruction process: the minimal BLAST alignment score and the maximum allowed nucleotide insertions between oligonucleotides.

For CE, data recovery relied on a de-Bruijn graph reconstruction approach, leveraging known header and tail oligos to identify the most frequent next oligo. CE required only one header and tail oligo, whereas CCE required two to achieve recoverability. De-Bruijn graph decoding is a common strategy in DNA storage applictions because it can reconstruct sequences even when individual fragments are missing or corrupted [45, 46]. Unlike traditional DNA strand recovery using de Bruijn graphs, we focus on chains of recovered oligos. By following statistically dominant transitions between overlapping oligo sequence segments. The algorithm tolerates sparser sequencing coverage and moderate error rates.

For PE an alternative decoding strategy was used. Instead of reconstructing complete strands, the algorithm detected positional triplets, locating pairs of positional motifs with precisely one data-encoding oligo in between. For each position, the most probable data oligo was selected. This approach requires only local identification of three-oligo combinations rather than full-strand reconstruction. Similar block-based decoding strategies have been explored in block DNA storage systems, where dividing sequences into smaller independently decodable units improves robustness to sequencing noise and strand loss [16]

All original images were successfully recovered under appropriate decoding parameters.

Recovery success for one purification step is shown in the heatmaps in Figure 4. These heatmaps show how decoding parameters influence recoverability. They indicate that a single purification step was generally sufficient to remove byproducts, enabling reconstruction of the expected sequence. Additional purification steps reduced nonspecific products but often removed too much of the target strand, preventing recovery because all assemblies were performed at the same oligonucleotide concentration. Two purification steps may further improve recovery only if sufficiently high oligonucleotide concentrations are used. Each assembly was done in duplicates.

**Figure 4.**
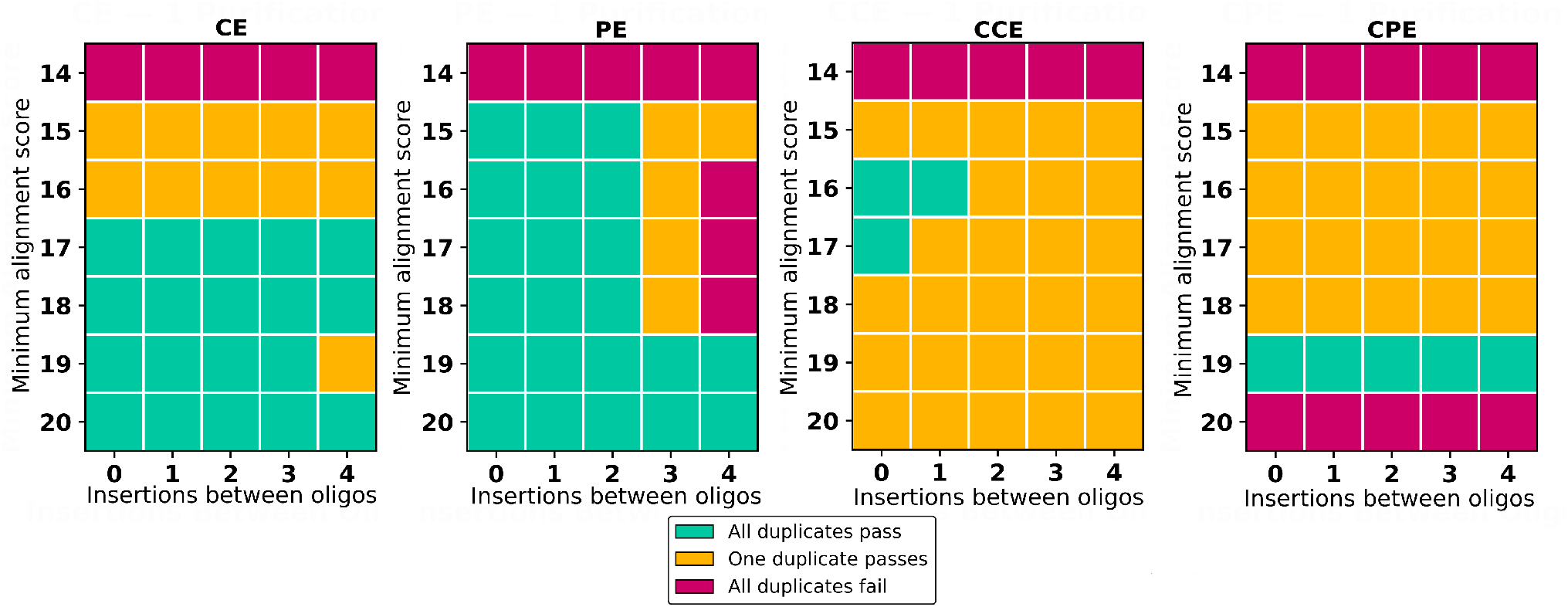
Recovery of assembled sequences from once-purified DNA for all four encoded images under varying parameters. The x-axis indicates the maximum allowed Insertions between two oligonucleotides during recovery, and the y-axis shows the minimum alignment score in the BLAST alignment. The color indicates the amount of duplicates that were able to be recovered.

#### Error Correction

Recovery was possible up to a minimal BLAST alignment score of 15 out of 20 (5 errors) regardless of the encoding. Our library’s edit distance of 3 per domain enables a theoretical error correction of 30% (6 errors / 20 nt oligo length). In principle, combined left and right domain distances allow error correction across both domains if errors are distributed evenly. In practice we achieved 25% correction because of unfavorable error distributions. This demonstrates the inherent error-correcting capability of the DNA block assembly approach independent of the encoding scheme. In contrast, *de novo* synthesis methods typically employ inner and outer error-correcting codes, applying the same principle of restricting sequence space. These methods generally achieve higher sequence density but lower inherent error correction. [5, 47, 36].

#### Error analysis of canonical encodings

We analyzed the assembly of canonical encodings by aligning recovered oligo chains to expected oligo chains (Figure 5a). The diagonal representing complete homology shows higher counts than partial matches. The clustering around the homology line indicates predominant correct assembly. Using the aligned recovered sequences, we determined where recovery failures occur (Figure 5b). Misalignments appeared around certain individual oligos. These oligos exhibits low ligation efficiency, hinting that some oligos block assembly of the final strands. Alignment dips also appear noticeably between fragments excised in the first purification set. Oligos which were added after the first ligation set exhibited the worst recovery rates.

**Figure 5.**
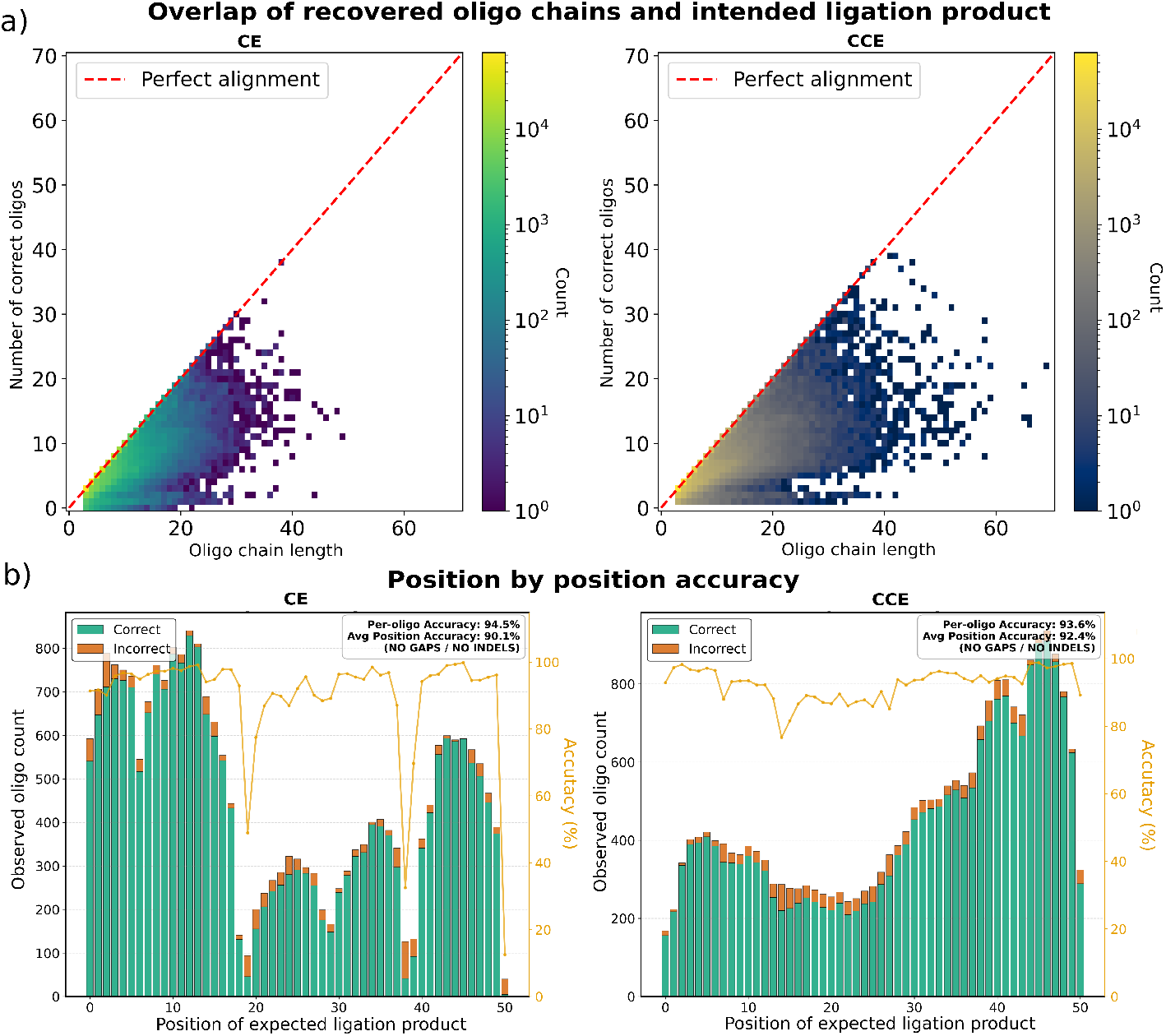
Sequence data analysis of recovered canonical encoding pictures. a) The heatmap depicts the number of correctly aligned oligonucleotides to the expected ligation product. Points located on the dotted red line represent sequences that are exact substrands of the expected oligo sequence. b) Count of correct and incorrect oligo alignments at each position of the expected ligation product, highlighting positional fidelity across the recovered data. (Maximal BLAST alignment score 20, Nucleotide insertions 0)

Canonical encodings was more resilient to sparser coverage, however proved more sensitive to incorrect hybridizations. Misassembled oligo sequences introduced incorrect edges in the de-Bruijn graph, effectively randomizing the graph and preventing reliable path reconstruction. CCE uses multiple strands together. Repeated oligos across these strands require larger headers and tails spanning two oligos to separate the 5 distinct strands. Even then, the de-Bruijn graph formed one linked graph. The recovery paths divert between strands and reduce overall success.

#### Error analysis of the positional encoding

Data recovery for PE proves more robust, because it only requires the retrieval of correct triplets. Since each triplet forms with just one ligation, decoding probability is higher compared to canonical encoding. However, without purification incorrect triplets can outnumber correct ones, while excessive purification causes high triplet loss. Both reduce recoverability. Larger fragments or higher combinatorial parameters thus require higher initial oligo concentrations or more accurate purification.

PE recovery searches for triplets with matching left and right motifs for all positions. Correct triplets must outnumber erroneous triplets sharing positional motifs. The ratios of the correct triplet to the highest erroneous triplet at a given position are illustrated in Figure 6a and Figure 6b for alignment score 20 with one purification. Lower ratios show background noise drowning out correct signals. One triplet in PE experiences exceptional noise. Positional motifs can coincidentally span two expected triplets. This in addition to a low ligation efficiency of the involved oligos may cause this noise. Cross-triplet detection and correction enables a higher recovery rate. The high cross-ligation potential of CPE increases background noise, leading to failure under the most restrictive recovery conditions.

**Figure 6.**
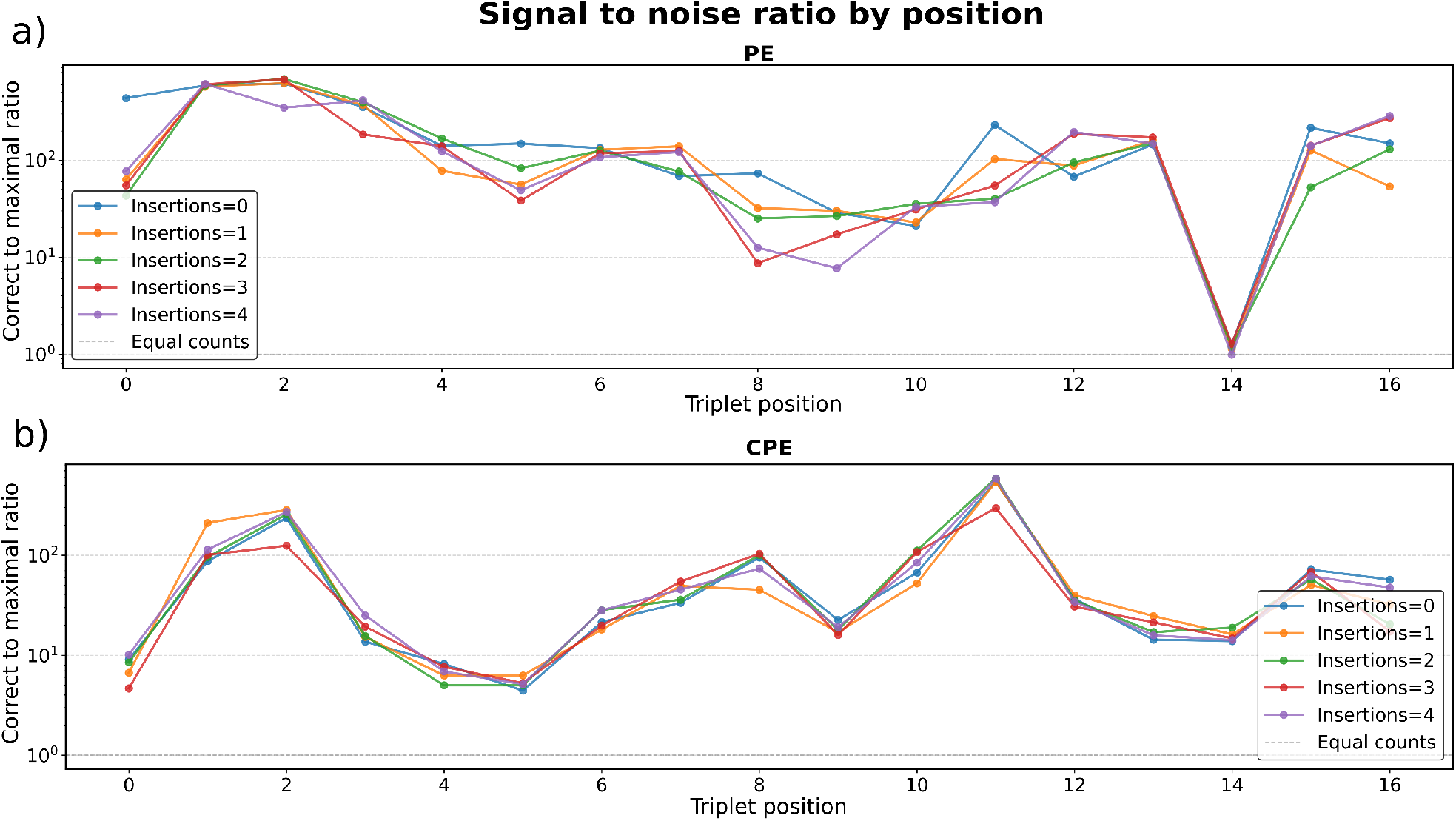
Sequence data analysis of triplet recovery in the positional encoding schemes. a) For PE, the ratio of the count of the expected triplet to the most abundant incorrect triplet at each position, using alignment scores of 20. b) For CPE, the ratio of the count of the expected triplet to the most abundant incorrect triplet at each position, using alignment scores of 20.

### Computing using staple libraries

We constructed toehold-mediated strand displacement gates consisting exclusively of library oligo concatenations to demonstrate the computational capability. These types of logic gates have been analyzed thoroughly before [48, 49]. Figure 7a shows gate structures and action pathways of the AND and OR gates. Supplementary Table S6 lists sequences for gate strands, input strands, and reporter strands.

**Figure 7.**
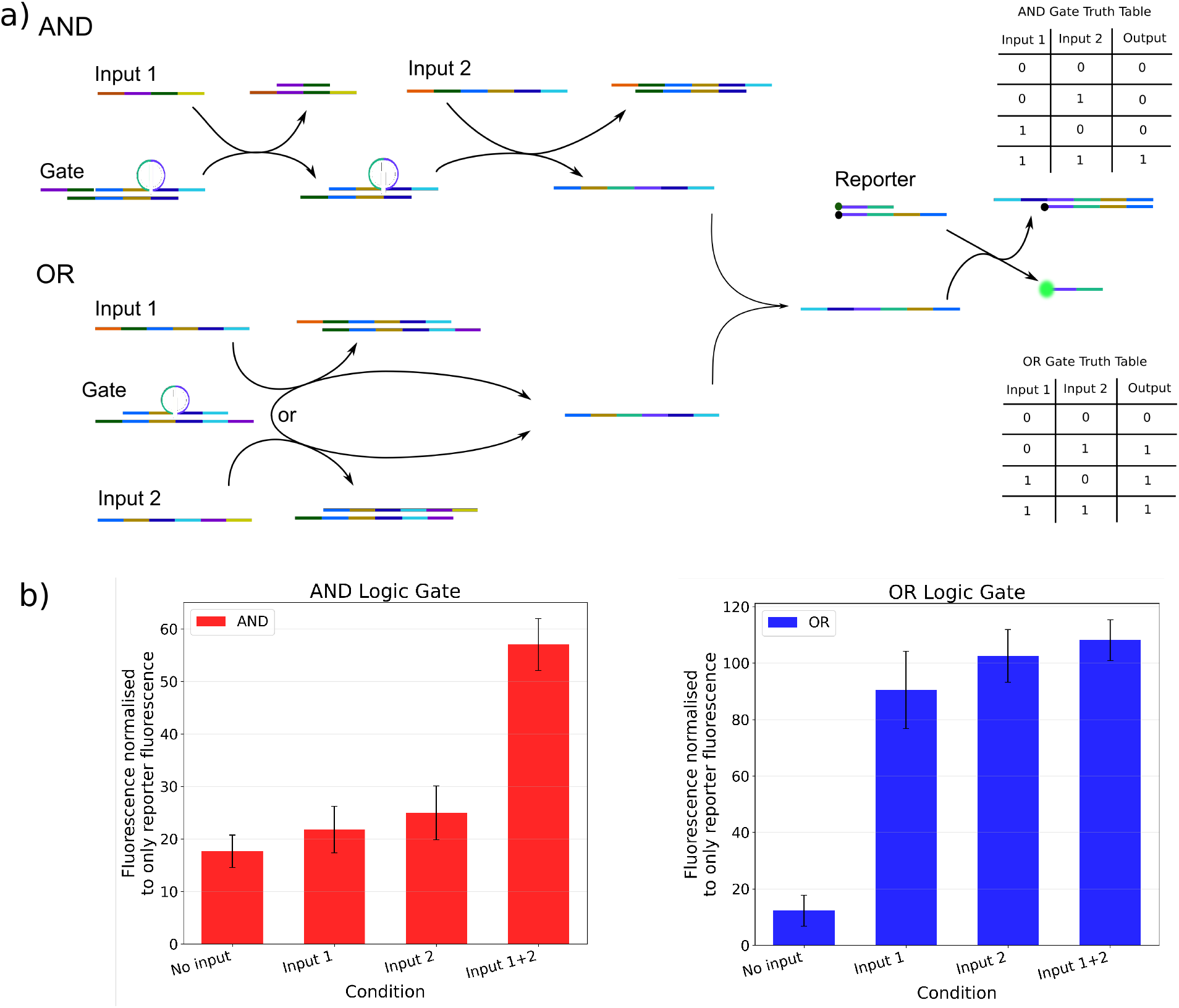
a) Strand displacement pathways for AND and OR gates. Colors indicate motifs from the DNA library, with two colors corresponding to each oligonucleotide. Therefore, all strands contain multiples of two motifs. Reverse complements are shown in identical colors. The corresponding truth tables for each gate are shown alongside the gates. b) Average fluorescence over eight replicates of the Gate–Reporter system for AND and OR gates under different input combinations.

Gates operate via toehold strand displacement, where input strands displace reporter strands from gate complexes. Reporter strands separate the fluorescein amidites (FAM) from Black Hole Quencher-1 (BHQ-1) in the reporter pair, producing measurable fluorescence. The average relative fluorescence of eight replicates is shown in Figure 7b. The truth table in Figure 7 confirms that both gates correctly implement their Boolean functions, with 0 denoting the absence of the input strand and 1 its presence. These gates, input strands, and reporters were assembled directly from salt purified DNA strands without additional purification. This shows that concatenations of library sequences can be used for both arbitrary data encoding and dynamic logic computation.

## Conclusion

The DNA staple library provides a dual-purpose platform for both DNA data storage and molecular computation. Unlike synthesis-heavy approaches, library elements can be mass-produced via bioreactors, eliminating costly *de novo* oligonucleotide synthesis. The use of a predefined oligonucleotide set shifts system complexity from synthesis to programmable assembly, allowing the same components to be used for multiple encoding and computational tasks.

A key advantage of the staple library approach lies in its potential to substantially reduce the cost of DNA-based data storage. In conventional synthesis-based DNA storage systems, each new dataset requires synthesizing large numbers of unique oligonucleotides, causing storage costs to scale directly with data volume. By contrast, the staple library is synthesized once, relies on desalted instead of HPLC-purified oligonucleotides, and can be used for arbitrary datasets. In this framework, the economic bottleneck shifts from sequence synthesis to programmable assembly and enzymatic processing, both of which can be automated and scaled using molecular biology workflows. This approach reduces the use of toxic chemicals and, due to its restricted sequence space, lowers potential biohazard risks. Consequently, stapling libraries provide a pathway toward significantly more economical DNA data storage systems, particularly for large-scale deployment.

The assembly of data-encoding strands is fully programmable and automatable. This enables calculations to occur directly on DNA encoded data avoiding the bottleneck of the Van-Neuman computing architecture and possibly enabling mass parallel computing. Our experiments demonstrate that different encoding schemes exhibit distinct trade-offs between data density, assembly reliability, and decoding robustness. In particular, canonical encodings provide the highest data density but show greater sensitivity to false hybridizations and assembly byproducts. In contrast, positional encodings achieve more consistent recovery by relying on local triplet identification rather than global strand reconstruction. These results highlight that encoding strategies must be co-designed with biochemical assembly protocols, as purification steps, ligation efficiency, and hybridization specificity influence data recoverability. The bioinformatic recovery of encoded data also depends on the chosen encoding scheme and can be optimized to further increase the probability of successful recovery. The flexible architecture of the staple library supports diverse encoding schemes optimized for assembly yield, automation, data density, or computational accuracy, thereby enabling application-specific customization across all stages of storage and processing. Designing encoding schemes that align with specific applications and computational tasks could give rise to specialized storage and molecular computing systems. This library likely enables broader applications, expanding computational ability beyond two-input Boolean functions to hybridization specific computing and, in specific cases, even NP-complete problems [50, 51, 52].

Data storage and Boolean computation was demonstrated with this library, however, broader applications demand a library redesign. Stronger energetic separation between intended and uninteded oligo interactions becomes essential, alongside new encoding schemes, protocols and recovery algorithms that maximize yield, specificity and recoverability. Incomplete nick ligation significantly limits the probability of assembling fully intact strands. Improvements in ligase efficiency would exponentially increase assembly yield and data recoverability.

While strand displacement gates are inherently leaky, our longer library strands compared to traditional approaches exacerbate erroneous displacement due to exposed 10 bp ssDNA overhangs on all interacting partners. This universal overhang design increases off-target toehold binding. A complementary library of reverse complement oligos could inhibit erroneous displacement by thermodynamically stabilizing correct gate complexes, providing a computational basis with significantly reduced leakage.

Our predefined oligo library circumvents DNA data storage’s primary scalability limitation, namely *de novo* synthesis costs, while simultaneously enabling seamless automation for high-throughput data writing and molecular computation at larger scales. By supporting both storage and computation within the same molecular framework, such libraries could form the foundation for scalable DNA-based information processing systems. With further advances in encoding design, reaction engineering, and automated assembly workflows, staple libraries may evolve into a universal platform for a universal molecular storage and programmable DNA computers.

## Experimental Section

### Assembly of stored data

Data-encoding strands were synthesized by first encoding information into library oligos, then partitioning into ligation pools containing only complementary motifs once, following precalculated pooling prescriptions. For primary pools, 50 pmol of each required oligo was combined with 1*/*4 Volume T4 ligation master mix which is a 1:2:2 ratio of T4 DNA ligase (26 U oligo^*−*1^, New England Biolabs, Germany, Cat. No. M0202), T4 ligase buffer (1*×* final concentration, New England Biolabs, Germany, Cat. No. B0202), DNase-free water. Each pool was incubated at 25°C for 30 min, then heat-inactivated at 65°C for 10 min. Non-primary pools were concentrated to 5 *µ*L prior to identical ligation using the entire volume.

### Purification of Ligation pools

Ligation pools were resolved on 4% (w/v) agarose gels prepared in TAE buffer (1 *×* final concentration, Thermo Fisher Scientific, USA, Cat. 15558026) and SYBR Safe DNA gel stain (0.006% (v/v), Thermo Fisher Scientific, Germany, Cat. No. S33102). Samples were loaded with DNA Loading Dye (1 *×* final concentration, Thermo Fisher Scientific, Germany, Cat. No. R0611). The electrophoresis was conducted at 4 V cm^*−*1^ for 3 h, and gels were imaged using a FastGene FAS-X system (Nippon Genetics, Germany) Bands corresponding to expected ligation products were excised and extracted using QIAquick Gel Extraction Kit (Qiagen, Germany, Cat. No. 28704) per manufacturer’s protocol (gel slices ≤ 400 mg solubilized in 6 *×* Volume Buffer QG at 50 °C, eluted in 30 *µ*L DNase-free water). Extracts were concentrated by ethanol precipitation: 0.1 *×* Volume 3 M sodium acetate (pH 5.2) and 2.5 *×* Volume ice-cold 100% ethanol added, incubated at *−*20 °C (30 min), pelleted (17000 *× g*, 30 min, 4 °C), supernatant removed, pellet washed with 500 *µ*L 70% ice-cold ethanol, repelleted (17000 *× g*, 30 min, 4 °C), supernatant aspirated, air-dried, and resuspended in 5 *µ*L DNase-free water.

### Sequencing of encoded data

Library preparation was performed using the Native Barcoding Kit 96 V14 (Oxford Nanopore Technologies, United Kingdom, Cat. No. SQK-NBD114.96) according to the manufacturer’s protocol. Pooled barcoded assemblies were sequenced on a MinION Mk1D using a FLO-MINSP6 flow cell (Oxford Nanopore Technologies, United Kingdom, Cat. No. FLO-MINSP6).

Raw data was basecalled with Guppy (dna_r9.4.1_-450bps_sup.cfg). We processed barcoded reads using the following process for our custom algorithm:

1. BLAST alignment of library elements to sequencing data (S3),
2. identification of library element chains (S4, 14 *≤ 𝓁*_*min*_ *≤* 20, 0 *≤ g*_*max*_ *≤* 4, *h*_*min*_ = 3) and
3. recover data from library element chains (PE and CPE: S6; CE and CCE: S7, *k* = 6, *β* = 200)

### Gate assembly and readout

Gate parts annealed together at equimolar concentrations (95°C → 16°C at 0.1°Cs^*−*1^) using a thermocycler (Eppendorf AG, Germany, Cat No. EP6381000018). Reporter strand where annealed together using a ratio of 1:1.2 of fluorescent stand to quencher strand. Gates (50 pmol), reporter (50 pmol), and inputs (50 pmol) were combined. No-input controls used 5 *µ*L DNase-free water for each missing input.

Fluorescence was monitored for 10 min at 37°C on a LightCycler 480 (Roche Diagnostics, Switzerland, Cat No. 05015278001) excitation wavelength 498 nm /emission wavelength 510 nm.

## Supporting Information

Supporting Information is available from the Wiley Online Library or from the author.

## Acknowledgements

This work was funded by the European Union (MI-DNA DISC, 101115215). Views and opinions expressed are however those of the authors only and do not necessarily reflect those of the European Union or the European Research Council Executive Agency. Neither the European Union nor the granting authority can be held responsible for them.

