## Supplemental information for "DNA Staples: An oligonucleotide library for data storage and computing"

**9 References** **33**

### Supplementary Note 1: Library analysis

Because the maximal density of a library $A$ is given by

$$d = \frac{\log_2(|A|)}{n}$$

it depends on both the library size $|A|$ and the oligonucleotide length $n$. The choice of $n$ strongly depends on the desired error-correction capability of the library. Figure S1a shows the average edit distance for motif sets of varying sizes and strand lengths. As expected, larger strand lengths and smaller domain sets increase the average edit distance.

The oligonucleotide length additionally influences the energetic separation between intended and unintended interactions. As oligonucleotide length increases, the sequence space grows exponentially, allowing the selection of motif sets with reduced unintended complementarity and therefore larger energetic separation between intended and unintended interactions. Figure S1b shows the distribution of energetic gap sizes for libraries generated with different oligonucleotide lengths. Larger oligonucleotides generally produce larger energetic gaps, enabling complete energetic separation at strand lengths of 36 nucleotides.

One caveat of this analysis is that NUPACK predicts an increasing number of configurations for longer strands. As a result, for unintended interactions the most likely configuration becomes less well defined. Nevertheless, the predicted energies still approximate the configuration with the strongest binding interaction.

To further illustrate density behavior in the encodings used in this work, Figure S2 presents density changes as a function of library size, oligonucleotide length, and combinatorial parameters.

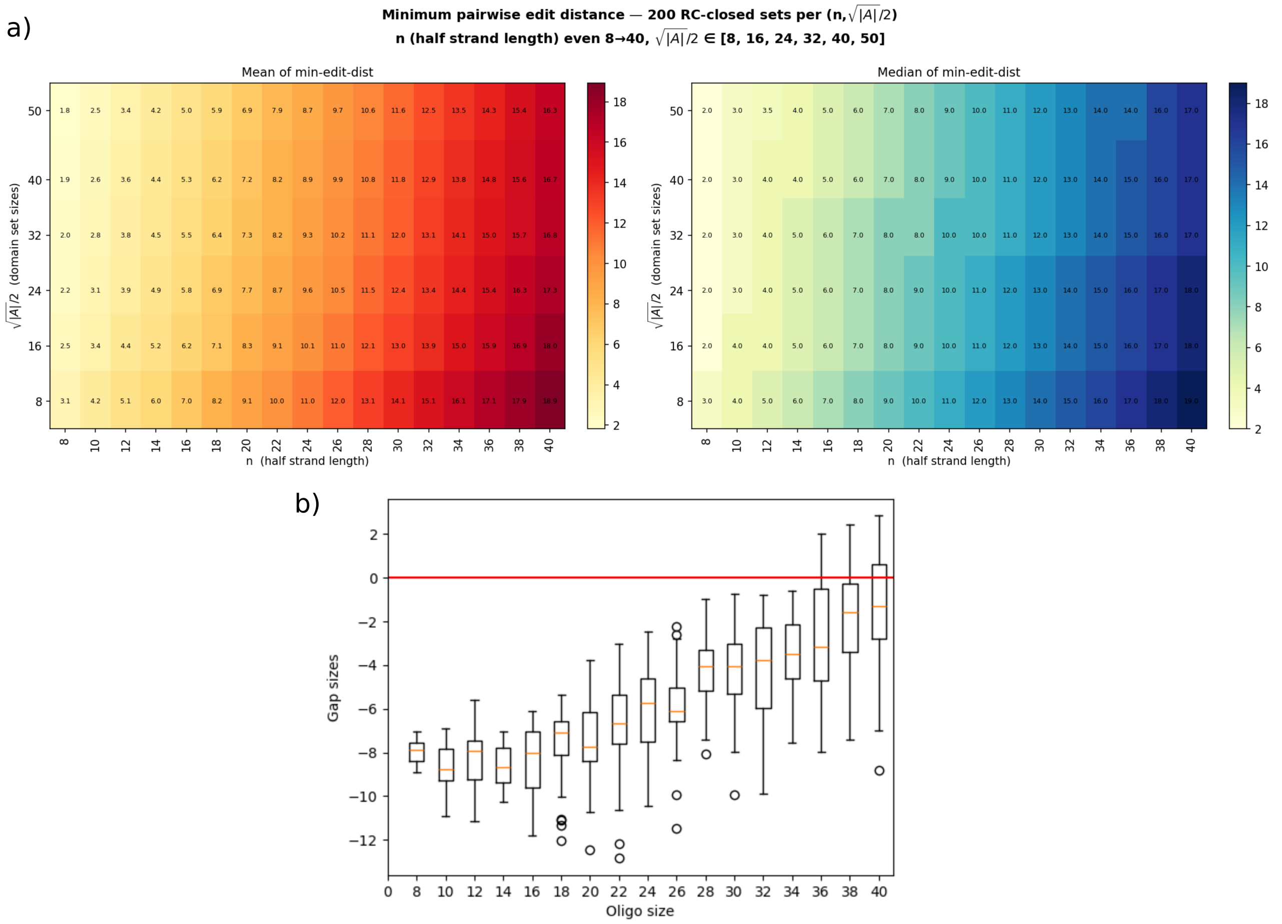

**Figure S1.** Considerations for selecting oligonucleotide lengths for the library. (a) Average and median minimum edit distances between domain motifs as a function of domain strand length and library size, computed over 200 randomly generated domain sets. (b) Binding energy gap sizes between intended and unintended interactions for 20 randomly generated libraries.

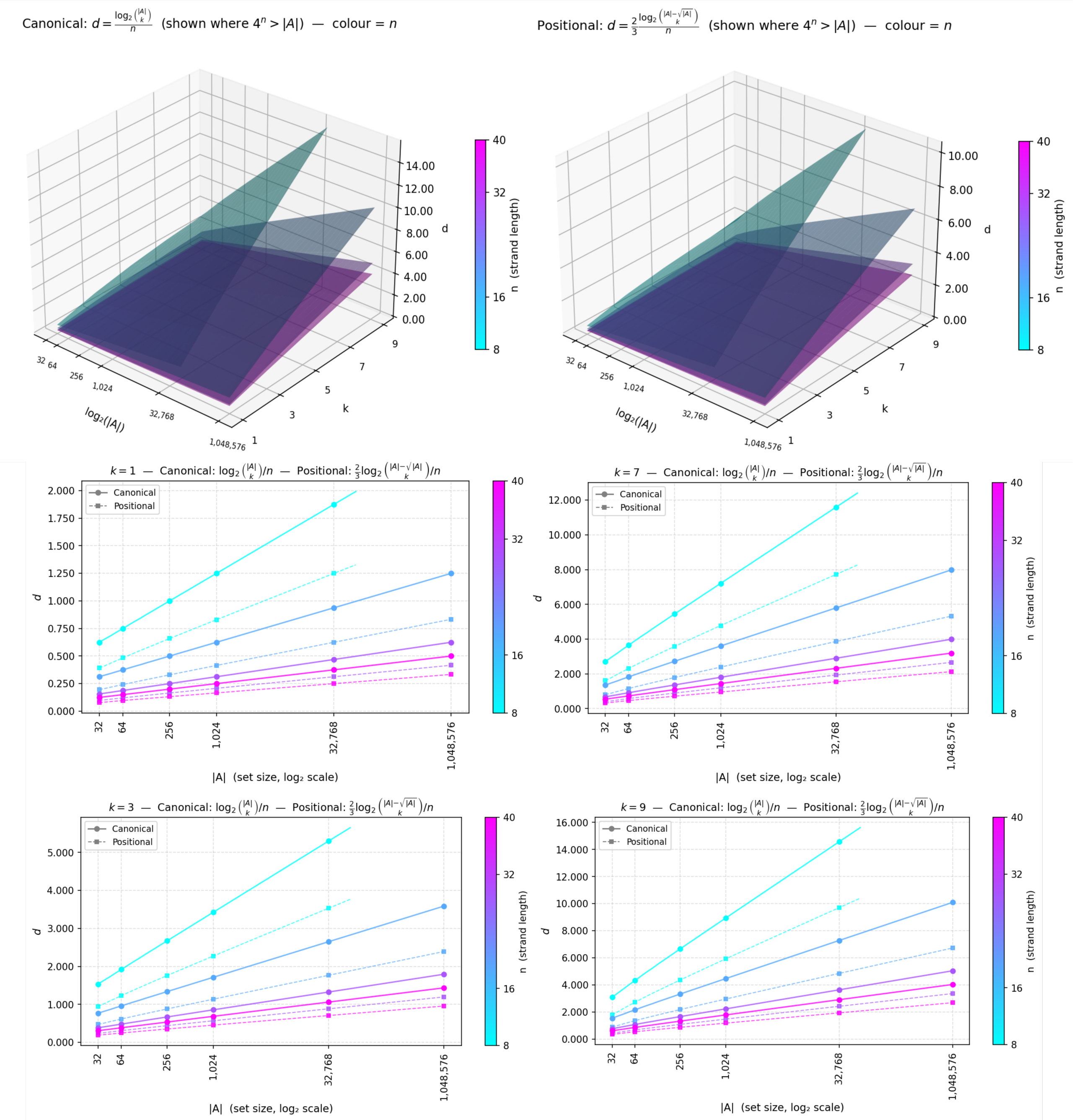

**Figure S2.** Densities of both canonical and pisitional encodings in depenedence of the library size, oligo length and combinatoprial factor $k$

### Supplementary Note 2: Assembly analsis

#### A. Gel electrophoresis

Here we show all agarose gels prior to purification (Figure S3).

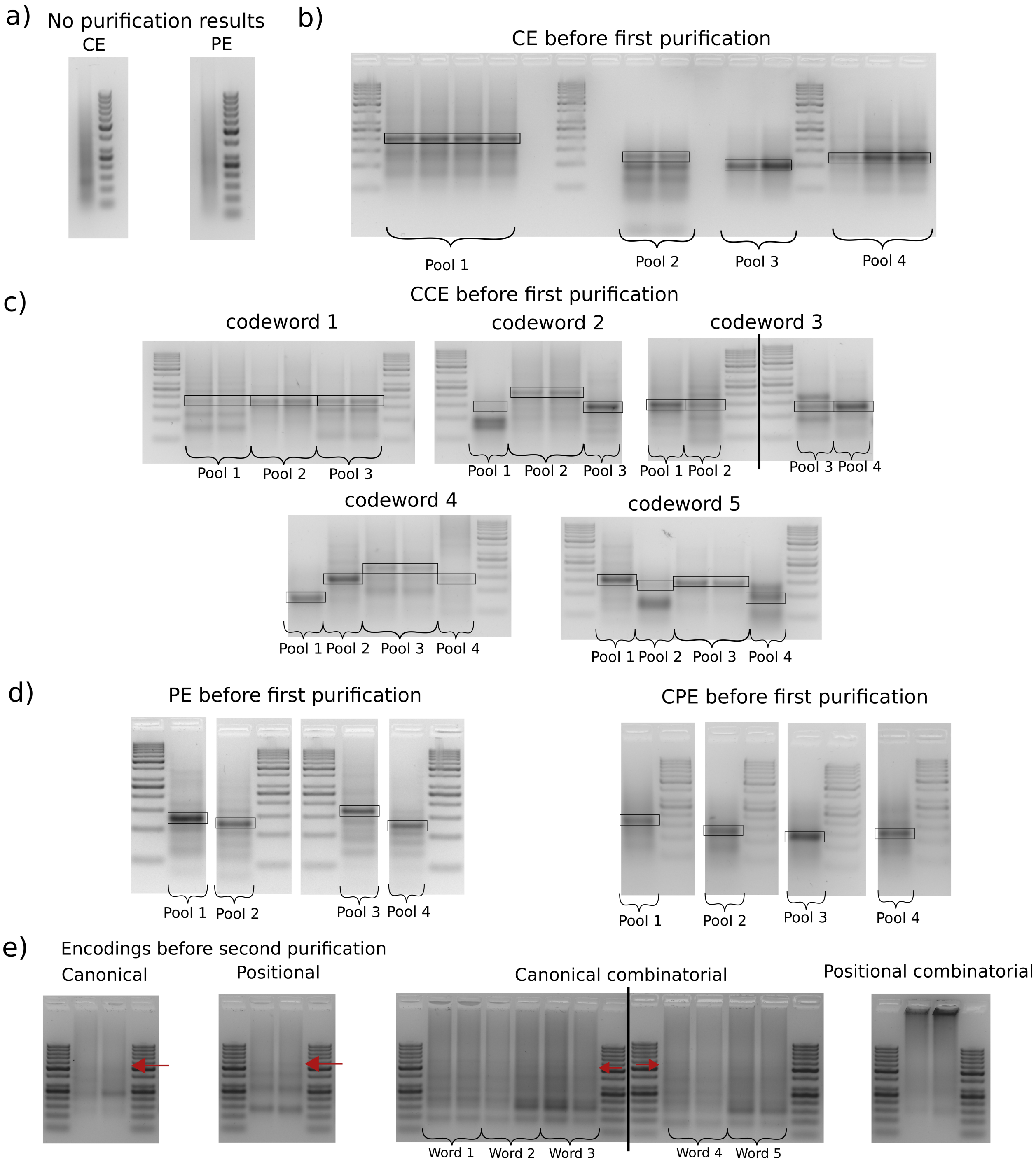

**Figure S3.** Gel electrophoresis analysis of strand assembly and purification. a) Assembly results obtained without intermediate purification for CE and PE b-d) Agarose gel of the all encodings prior to the first purification step. Boxes indicate the expected target length for each pool. e) Agarose gel showing the four encoded images after final assembly and prior to the second and final purification step. All codewords are present in duplicate. Red arrows indicate the target band size used for purification where applicable.

#### B. Oligo chain recovery

After applying oligonucleotide chain recovery, we analyzed what fraction of base-called strands can be mapped back to oligonucleotide chains. This depends on the chosen recovery parameters. All recoveries were performed with a minimum chain length of 3, which discards all chains shorter than this threshold. This parameter itself affects recovery success: a higher minimum chain length can filter out random oligonucleotide ligation events but may also remove valid information, whereas a lower threshold increases noise.

Under the least restrictive recovery parameters (minimum BLAST alignment score 14, insertions between oligos = 4), on average 42.7%–52.0% of base-called strands are remapped to oligonucleotide chains. In contrast, the most restrictive parameters yield an average recovery of only 29.1%–34.2% (Figure S4).

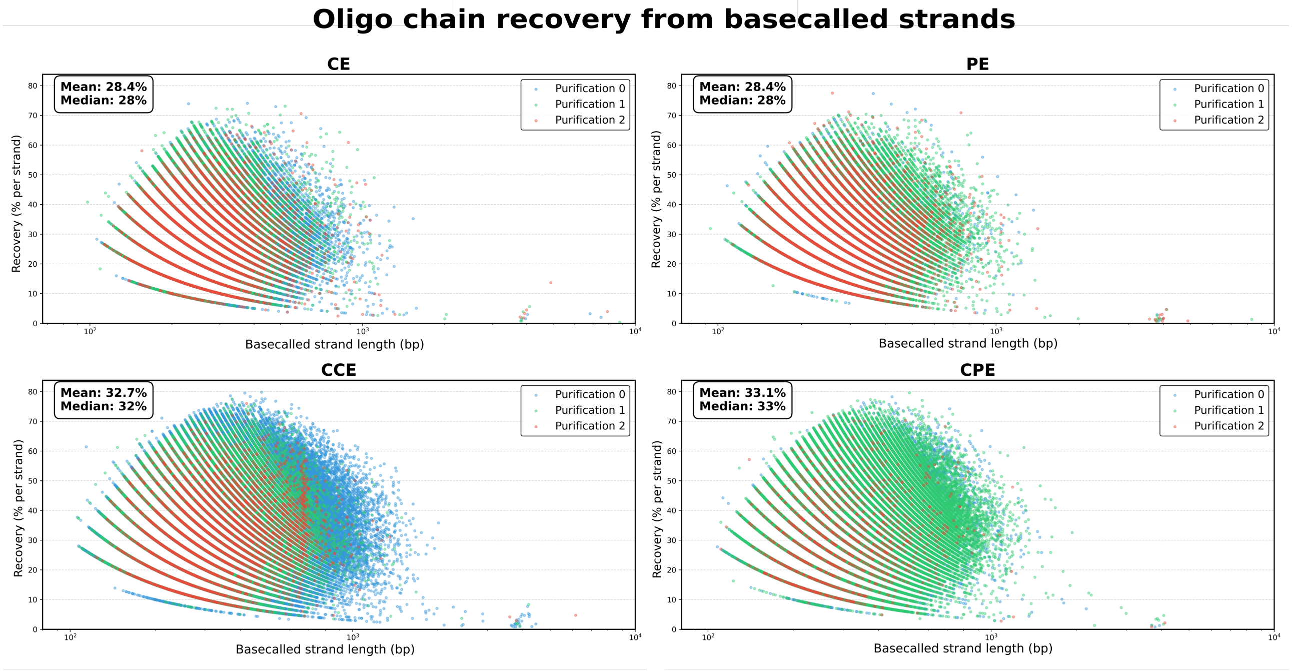

**Figure S4.** The heatmap depicts the number of correctly aligned oligonucleotides to the expected ligation product seperated by purification level. Points located on the dotted red line represent sequences that are exact substrands of the expected oligo sequence.

#### C. Alignment analysis

For correct assembly analysis as presented in this paper, a strict alignment procedure was used (Supplementary Algorithm 5), allowing only shifts and substitutions without gaps or insertions. This is applied to the oligonucleotide chain derived directly from the BLAST alignment. This strand is used for the de Bruijn graph reconstruction, where each error propagates as an incorrect edge in the graph.

While we previously showed the overlap of correct oligonucleotides with the intended ligation product across all purification conditions combined, here we additionally present the overlap analysis separated by purification level (Figure S5). Interestingly, samples with two purification steps show a pronounced spike at low alignment scores. This is likely due to reduced sample input in these conditions, where misligations of similar fragment sizes become more prominent (Section A).

We also present the per-codeword breakdown of alignment errors for each purification condition (Figure S6).

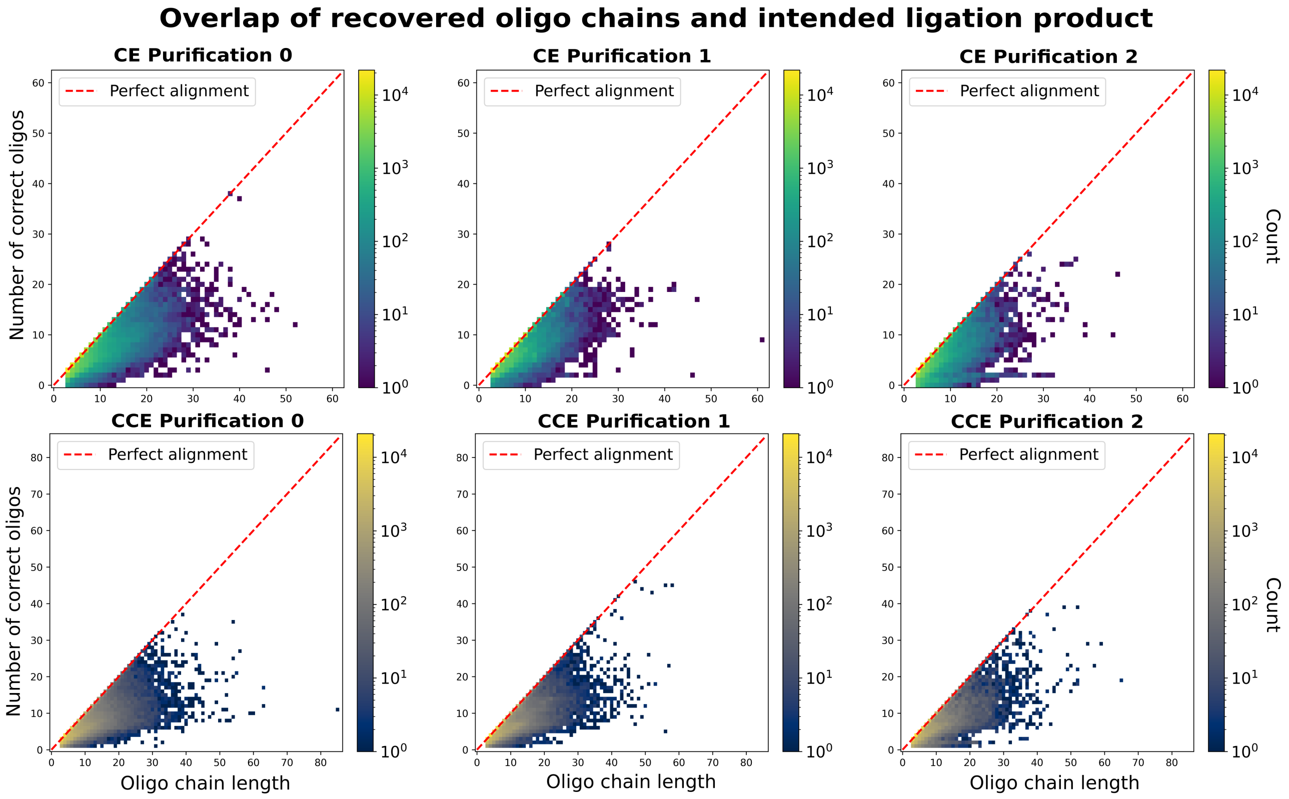

**Figure S5.** The heatmap depicts the number of correctly aligned oligonucleotides to the expected ligation product seperated by purification level. Points located on the dotted red line represent sequences that are exact substrands of the expected oligo sequence.

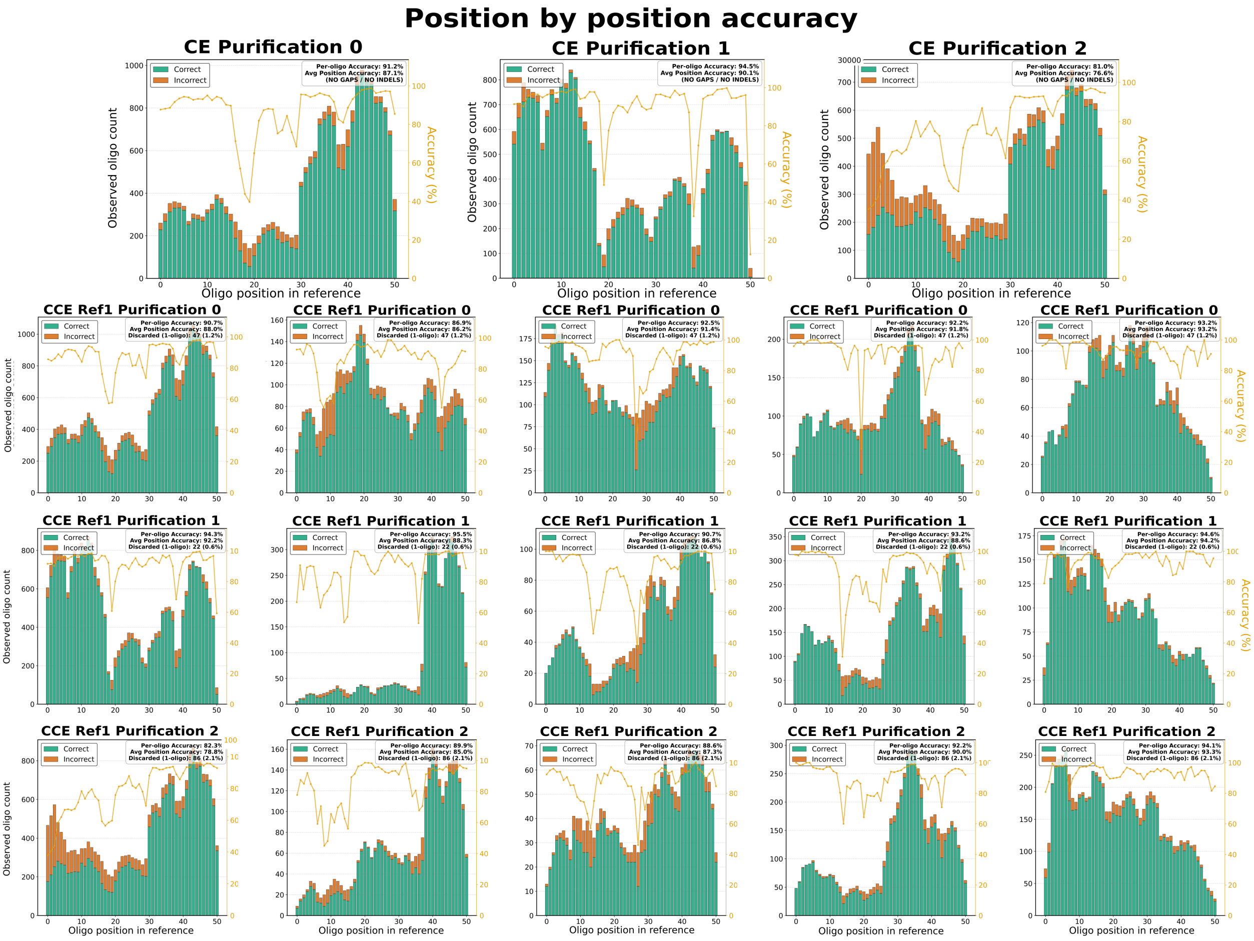

**Figure S6.** All alignment errors for all purification and intended assembly products for minimum BLAST alignment 20 and insertions between oligos 0

### Supplementary Note 3: Recovery analysis

#### A. Recovery of other purification levels

While we previously showed recovery success for a single purification, which yielded the most recoverable condition, other purification levels were also analyzed. Without purification, ligation products were severely smeared and no recovery was possible (Figure S7).

In contrast, two purification steps suffer from reduced coverage, as all assemblies were performed at the same initial oligonucleotide concentration. Under these conditions, CPE shows substantial cross-hybridization and smearing during the second purification, leading to significant product loss. In contrast, PE assembles in a more deterministic manner and remains recoverable. CE and CCE, however, are not recoverable, likely due to misligation (Figure S8).

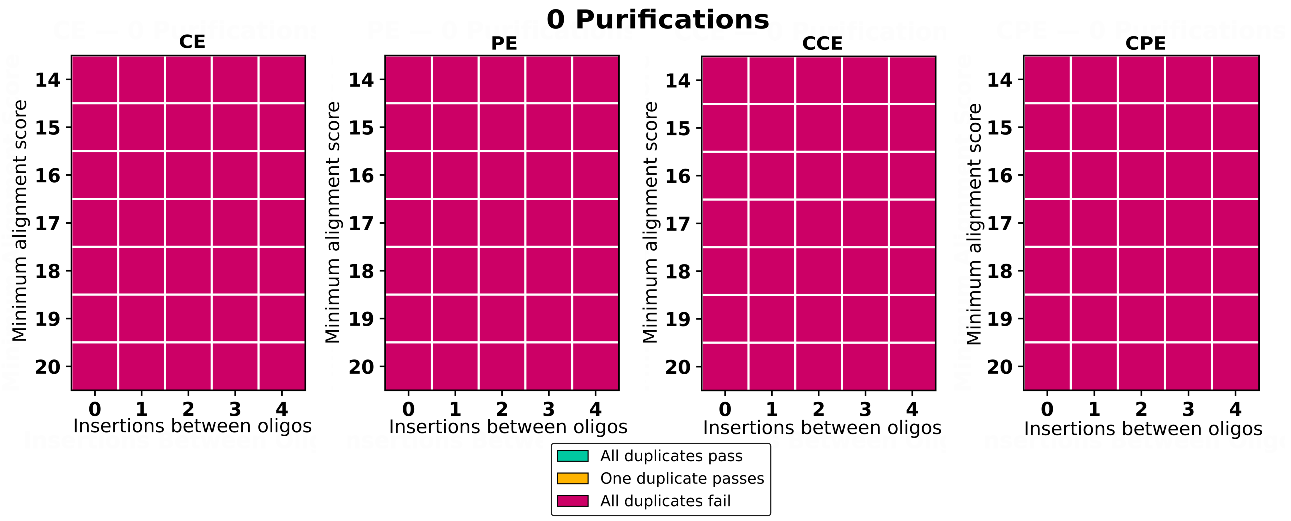

**Figure S7.** Recovery of assembled sequences from nonpurified DNA for all four encoded images under varying parameters. The x-axis indicates the maximum allowed Insertions between two oligonucleotides during recovery, and the y-axis shows the minimum alignment score in the BLAST alignment. The color indicates the amount of duplicates that were able to be recovered.

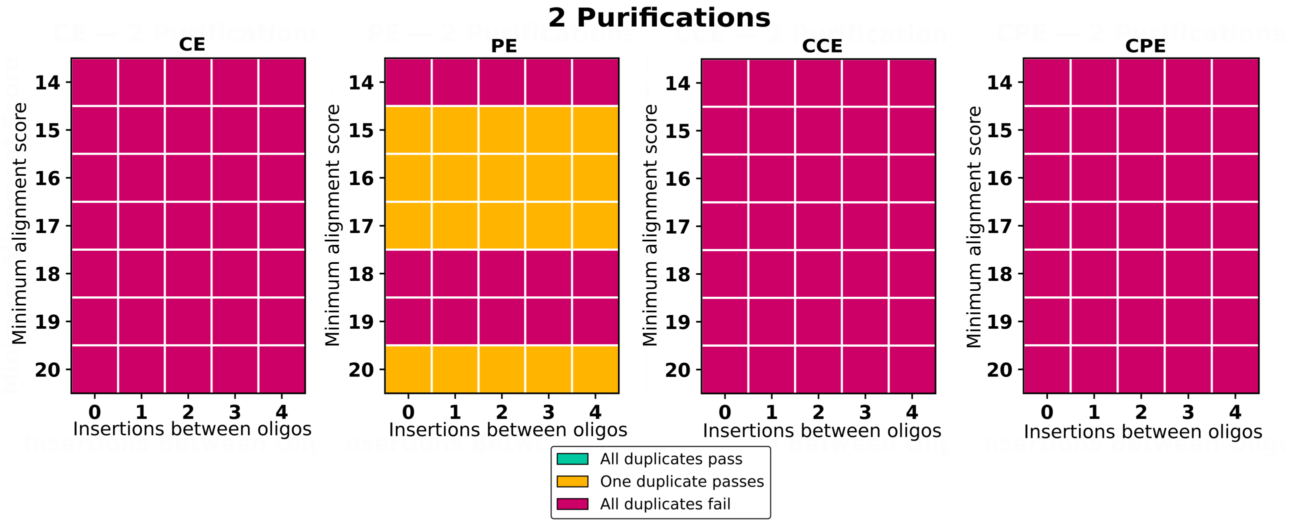

**Figure S8.** Recovery of assembled sequences from twice-purified DNA for all four encoded images under varying parameters. The x-axis indicates the maximum allowed Insertions between two oligonucleotides during recovery, and the y-axis shows the minimum alignment score in the BLAST alignment. The color indicates the amount of duplicates that were able to be recovered.

#### B. Blunting everything

We have shown that blunting only end motifs leads to recovery issues, as unligated strands introduce erroneous alignments, loss of information, and truncation of de Bruijn graph edges. To analyze this effect, we sequenced the same replicates while adding all library motifs to the samples, effectively maximizing blunting (Figure S9).

For the no-purification condition, this approach yields more recoverable information in PE and CPE, as additional triplets are blunted and fully sequenced even when they are freely floating and not bound to neighboring oligonucleotides. This enables partial recovery of both PE and CPE. CE and CCE, however, remain affected by their inherent misassembly sensitivity. Similarly, very sparse samples observed in the twice-purified condition are recovered more abundantly, with overlapping strands becoming more common, which also makes recovery of CE and CCE partially possible.
In contrast, under one-purification conditions, recovery performance decreases compared to specific motif blunting. This is likely because assemblies are relatively successful in this regime, resulting in a higher fraction of correctly ligated strands. Consequently, adding all motifs primarily increases background noise from misassembled strands and reduces overall recoverability. These results highlight that recovery success is governed by an interplay between yield, encoding strategy, and sample preparation.

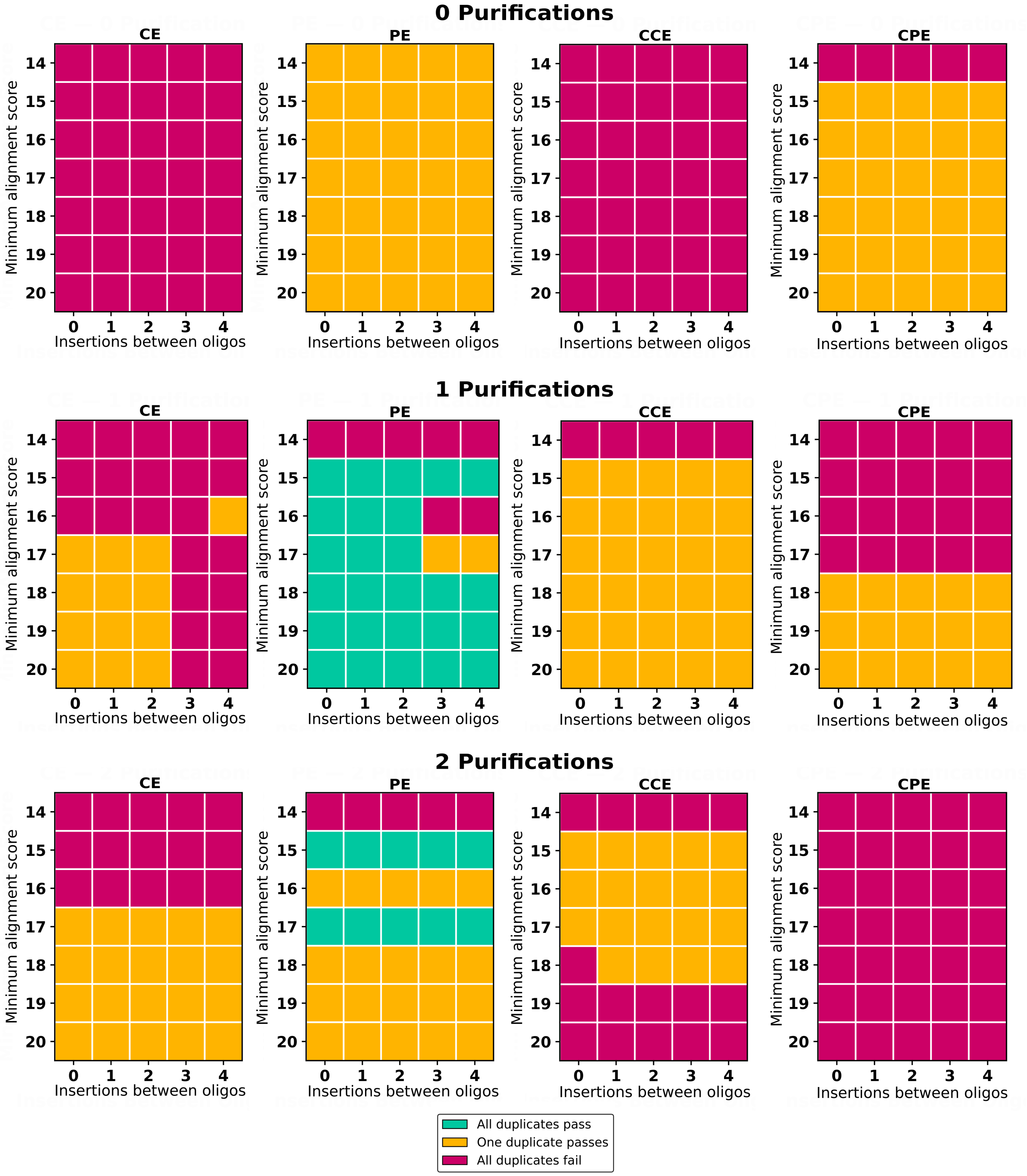

**Figure S9.** Recovery of assembled sequences from completly blunted DNA for all four encoded images under varying parameters. The x-axis indicates the maximum allowed Insertions between two oligonucleotides during recovery, and the y-axis shows the minimum alignment score in the BLAST alignment. The color indicates the amount of duplicates that were able to be recovered.

### Supplementary Note 4: Error analysis

#### A. Chimeras

Miss assembly effects are further amplified in CCE, as this encoding stores five intended ligation products simultaneously. Unligated fragments can therefore form chimeras between two different intended ligation products. To identify these events, we apply a filtering procedure (Figure S10). This filter removes all strands that exhibit at least two valid alignments, where one alignment transitions to another with at least two oligonucleotides per alignment segment. From the total 139,687 reads 37,504 are chimeric in this analysis for all recovery parameters. This shows that chimeras are a significant source of error.
In total, depending on purification conditions, between 80% and 88% of oligo chains in CCE are classified as non-chimeric for the tightest recovery parameter (BLAST alignment score 20, Insertions between oligos 0, purification 1: 3,991 oligo chains, purification 2: 3,985 oligo chains, purification 1: 3,937 oligo chains) (Figure S11). Chimeras may also contain more than one switch between intended ligation products, however, more complex chimeric structures become increasingly unlikely. These biases propagate into the assembly graph as uneven edge coverage and local disruptions in reconstructed ligation paths, ultimately affecting recoverability.

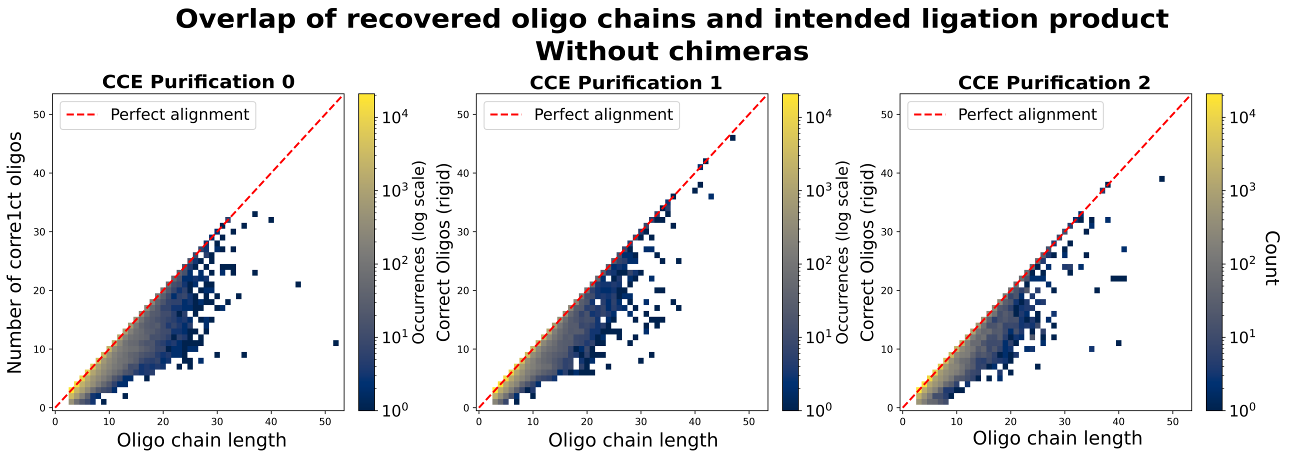

**Figure S10.** The heatmap depicts the number of correctly aligned oligonucleotides to the expected ligation product for CCE when all chimeres are removed. Points located on the dotted red line represent sequences that are exact substrands of the expected oligo sequence.

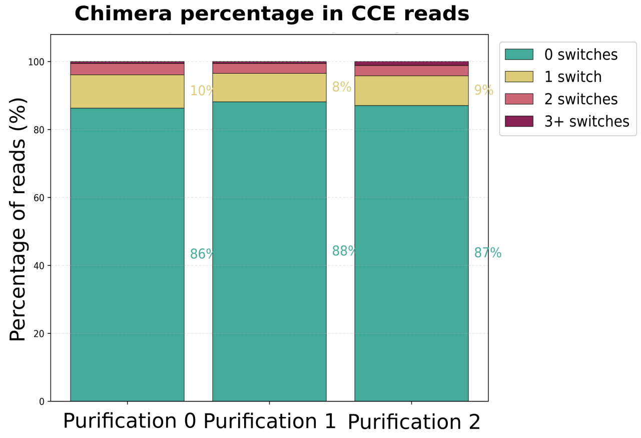

**Figure S11.** Percentage of chimeras percent in each purification for CCE. a chimera has at least 2 oligos of two different intended ligation production adjacent to each other in a read

#### B. Ligation errors

While chimeras fall under ligation errors, we further analyze recovered oligonucleotide chains to identify where in the ligation process these errors occur (Figure S12). This reveals that certain oligonucleotides exhibit lower ligation efficiency. While some peaks correlate with pools that fail to ligate, other peaks indicate that specific oligonucleotides themselves have reduced ligation propensity. These effects reflect both pool-level and sequence-intrinsic biases in ligation efficiency that propagate into assembly errors.

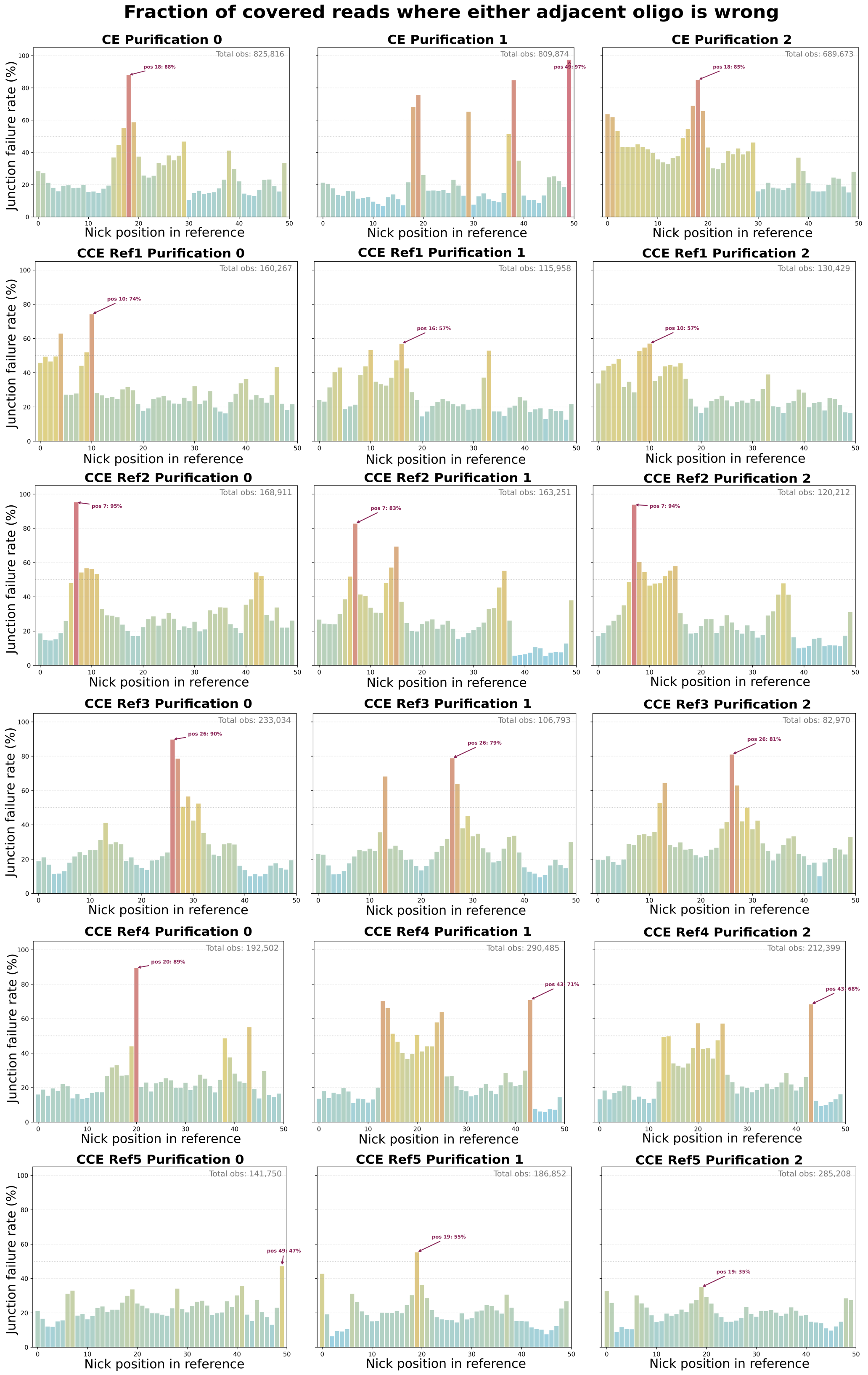

**Figure S12.** Per-position error counts of misliagtions for all pruificaitons and inted ligation products (Refrences) for minimum VLAST alignment score 10 and zero insetions between oligos. An error oocur escactly when an aligned strand at two naibouring positions $(i, i+1)$ the correct alignment

#### C. Error type analysis

So far, alignment analysis was performed using a rigid scheme that does not allow gaps and only accounts for substitutions. This choice was motivated by the use of a de Bruijn graph constructed from k-mers, where insertions or deletions would introduce incorrect graph edges. To further analyze the types of errors occurring in the recovered oligonucleotide chains, we implemented a semi-global Needleman–Wunsch algorithm [1]. The semi-global formulation allows alignments that do not necessarily start or end at the boundaries of the reference sequence, preventing artificial inflation of insertions caused by forced end-to-end alignment. From this alignment, we classify substitution, insertion, and deletion events at the oligonucleotide level (Figure S13, S14).

While substitution remains the dominant error type, insertions and deletions are also observed. These errors can arise from false hybridization events. Considering potential bioinformatic effects in the reconstruction process, the observed correlation between insertions and higher error tolerance in the oligonucleotides may indicate the presence of highly corrupted oligos within the assembly. These sequences may not be recognized as part of the intended oligonucleotide chains but, due to accumulated mutations, can be misattributed to incorrect library oligos during alignment. More flexible chain reconstruction may further allow strands containing multiple deletions within a single oligonucleotide to still be classified as continuous chains.

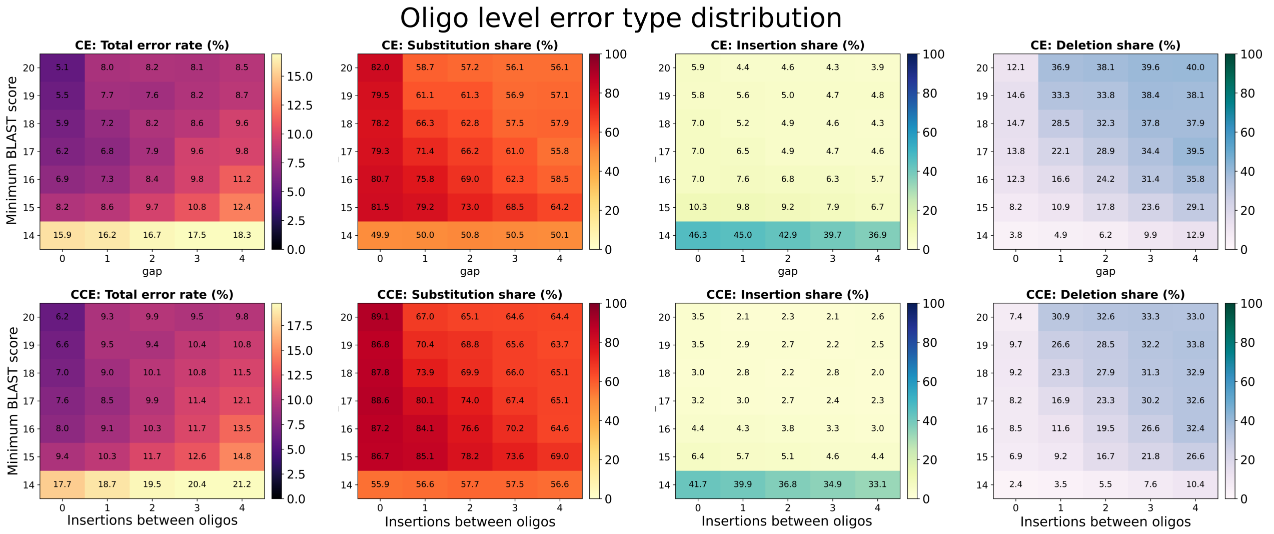

**Figure S13.** Error analysis across all recovery parameters. The first heat map shows the total error percentage, while the remaining heat maps show the distribution of error types, over substitutions, deletions, and insertions

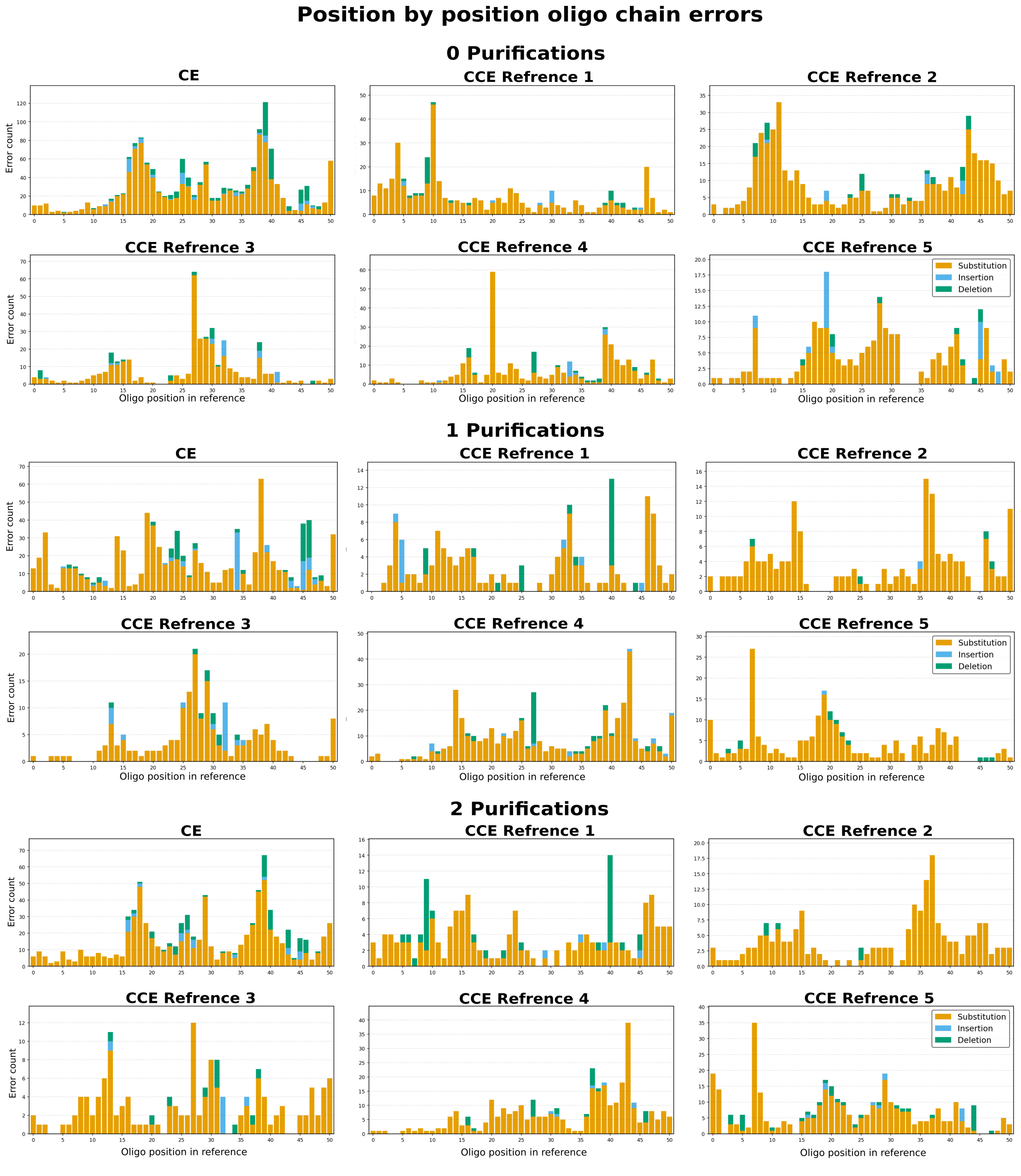

**Figure S14.** Per-position error breakdown across all intended ligation products (references), showing error counts and error types for recovery parameters with a minimum BLAST alignment score of 20 and zero insertions between oligonucleotides.

### Supplementary Note 5: Codes

In this section, pseudocode for the algorithms used in this paper is presented to provide an overview of how the storage and recovery pipeline was implemented.

#### A. Encoding

**Algorithm 1:** Canonical Encoding

**Input:** Binary data $D$, DNA library $\mathcal{L} = \{l_1, \dots, l_N\}$, barcode length $b$
**Output:** Set of DNA codewords $\mathcal{C}$
Sort $\mathcal{L}$ lexicographically;
Partition $D$ into binary codewords $\mathbf{W} = [w_1, \dots, w_m]$;
Convert each $w_i$ to decimal codeword $d_i = [d_{i,1}, \dots, d_{i,k}]$;
**foreach** *codeword* $d_i \in \mathbf{W}$ **do**
    $\mathbf{B}_i \leftarrow$ Counter$(b, N-1, i)$ ; `// Barcode indices`
    $c_i \leftarrow []$;
    **for** $j = 1$ **to** $k + b - 1$ **do**
        **if** $j \leq b$ **then**
            Append $\mathcal{L}[\mathbf{B}_i[j]]$ to $c_i$;
            $\text{left} \leftarrow \overline{\mathcal{L}[\mathbf{B}_i[j]][1:n/2]}$;
            $\text{right} \leftarrow \overline{\mathcal{L}[\mathbf{B}_i[j]][n/2+1:n]}$;
        **else**
            Append $\mathcal{L}[d_{i,j-b}]$ to $c_i$;
            $\text{left} \leftarrow \overline{\mathcal{L}[d_{i,j-b}][1:n/2]}$;
            $\text{right} \leftarrow \overline{\mathcal{L}[d_{i,j-b}][n/2+1:n]}$;
        Append $\text{left} \,\|\, \text{right}$ to $c_i$;
    Append final oligo $\mathcal{L}[d_{i,k}]$ to $c_i$;
    $\mathcal{C} \leftarrow \mathcal{C} \cup \{c_i\}$;
**return** $\mathcal{C}$;

**Algorithm 2:** Positional Encoding

**Input:** Binary data $D$, oligo library $\mathcal{L}$ with left motifs $M_L$ and right motifs $M_R$, barcode length $b$, selection size $k$
**Output:** Set of oligo-trio codewords $\mathcal{C}$

```
Construct positional libraries P_0, ..., P_{|M_L|+|M_R|-2} from L;
Partition D into binary codewords W = [w_1, ..., w_m];
foreach codeword w_i ∈ W do
    for each position p in w_i do
        Build candidate set S_p from Cartesian product of left/right motifs in P_{p+b};
        v ← int(w_i[p]);
        Initialize indicator vector z with first k entries set to 1;
        offset ← 0;
        while v ≥ offset + 2^u do
            offset ← offset + 2^u;
            Advance z to next k-combination in colex order;
        Select oligos O_p ⊂ S_p where z[j] = 1;
        r ← v − offset;
        f ← binary(r, u);
        foreach half-motif h_j ∈ O_p do
            if f[j] = 1 then
                replace h_j with \overline{h_j};
    B_i ← Counter(b, |P_0| − 1, i);
    for each position j in barcode + data do
        foreach oligo o at position j do
            if j is even then
                left ← M_L[⌊j/2⌋] ∥ \overline{o[n/2+1 : n]};
                right ← \overline{o[1 : n/2]} ∥ M_R[⌊j/2⌋];
            else
                left ← \overline{o[1 : n/2]} ∥ \overline{M_R[⌈j/2⌉]};
                right ← \overline{M_L[⌊j/2⌋]} ∥ \overline{o[n/2+1 : n]};
            Store trio (left, o, right);
    Add codeword to C;
return C;
```

#### B. Alignment

**Algorithm 3:** Barcode-wise Sequence Processing and Alignment against Oligo Library

**Input:** Set of barcode-specific sequencing folders, oligo library database $\mathcal{D}$
**Output:** Alignment files mapping sequences to library elements

```
foreach barcode group b in input dataset do
    collect all sequencing reads R_b from b;
    merge R_b into a single read set R_b^*;
foreach merged read set R_b^* do
    convert nucleotide reads into FASTA representation F_b;
foreach FASTA file F_b do
    align F_b against oligo library database D;
    store alignment results A_b in output collection;
return All alignment sets {A_b} indexed by barcode
```

**Algorithm 4:** Subsequence Extraction from Alignment Hits

```
Input: Alignment set B, minimum alignment length ℓ_min, maximum gap g_max, minimum hits per subsequence
       h_min
Output: Set of subsequences with associated query regions and hit collections
// Phase 1: Filter and organize alignment hits by query
H ← empty map from query identifiers to lists of hits;
foreach alignment record r ∈ B do
    extract query identifier q, subject identifier s, alignment length ℓ, and query-aligned interval (a,b);
    if ℓ < ℓ_min then
        continue
    (start, end) ← ordered interval of (a,b);
    append (start, end, s) to H[q];
// Phase 2: Build contiguous subsequences of hits
O ← ∅;
foreach query q in H do
    sort H[q] by start position;
    initialize empty subsequence C;
    initialize current interval bounds (s_start, s_end);
    initialize previous endpoint p ← undefined;
    foreach hit (start, end, id) in H[q] do
        if C is empty or (start − p) ≤ g_max then
            append id to C;
            update (s_start, s_end);
        else
            if |C| ≥ h_min then
                add (q, C, s_start, s_end) to O;
            C ← {id};
            reset interval to (start, end);
        update p ← end;
    if |C| ≥ h_min then
        add (q, C, s_start, s_end) to O;
return O;
```

**Algorithm 5:** Rigid (No-Gap Best-Fit) Alignment

**Input:** Read oligo sequence $\mathbf{r} = (r_1, \ldots, r_n)$, set of reference chains $\mathcal{C} = \{C_1, \ldots, C_k\}$, each $C_j = (c_1^j, \ldots, c_M^j)$
**Output:** Best-matching reference index $j^*$, offset $\delta^*$, match count $m^*$, per-position match/mismatch profile

```
m* ← 0;   j* ← ∅;   δ* ← 0;
// Orient read:  try forward and reverse complement
foreach orientation r′ ∈ {r, r̄} do
    foreach reference chain C_j ∈ C do
        // Try all integer offsets δ where read and reference share ≥ 1 position
        for δ ← −(n−1) to M−1 do
            i_start ← max(0, −δ);
            i_end ← min(n, M−δ);
            m ← 0;
            for i ← i_start to i_end − 1 do
                if r′_i = c^j_{i+δ} then m ← m+1;
            if m > m* then
                m* ← m;   j* ← j;   δ* ← δ;   r* ← r′;
// Record per-position match/mismatch against best reference
for i ← 0 to n−1 do
    p ← δ* + i;
    if 0 ≤ p < M then
        if r*_i = c^{j*}_p then match[p] += 1;
        else mismatch[p] += 1;
return j*, δ*, m*, match[·], mismatch[·];
```

##### C. Decoding

**Algorithm 6:** Triplet Alignment Recovery from Subsequence Data

**Input:** Subsequence file $\mathcal{S}$, well-to-motif map $\mathcal{W}$ : well $\to (m_L, m_R)$, left motif list $M_L$, right motif list $M_R$, minimum count $\mu$

**Output:** Position-indexed triplet map $\mathcal{T}$

Parse $\mathcal{S}$ into fragment lists $\mathcal{F} = [f_1, \ldots, f_N]$;

`// Step 1: Collect candidate triplets per motif-pair position`

**for** $i = 0$ **to** $|M_L| - 1$ **do**
- $m_L^{(i)} \leftarrow M_L[(i+1) \bmod |M_L|]$;
- $m_R^{(i)} \leftarrow M_R[i \bmod |M_R|]$;
- **foreach** *fragment* $f \in \mathcal{F}$ **do**
  - **for** $j = 0$ **to** $|f| - 3$ **do**
    - $w \leftarrow f[j..j+3]$;
    - $\mathbf{m} \leftarrow \big(\mathcal{W}[w_1], \mathcal{W}[w_2], \mathcal{W}[w_3]\big)$ flattened;
    - Detect left motif $\ell$ and right motif $r$ from $\mathbf{m}$;
    - **if** *both* $\ell$ *and its complement appear* ***or*** *both* $r$ *and its complement appear* **then continue** ;
    - $\mathbf{m}_c \leftarrow \min(\mathbf{m}, \mathrm{pairReverse}(\mathbf{m}))$;
    - **if** $\ell = m_L^{(i)}$ ***and*** $r = m_R^{(i)}$ **then**
      - $\mathrm{count}[\mathbf{m}_c] \leftarrow \mathrm{count}[\mathbf{m}_c] + 1$;
- $C_i \leftarrow$ top-5 canonical triplets in position $i$ with count $\geq \mu$;

`// Step 2: Pick highest-count candidate per position`

**foreach** *position* $i$ **do**
- Sort $C_i$ by count descending;
- $\mathcal{T}[i] \leftarrow C_i[0]$;

`// Step 3: Remove redundant triplets`

**foreach** *position* $i$ **do**
- $\mathcal{O} \leftarrow \bigcup_{j \neq i} \{\mathrm{wells}(t) : t \in C_j\}$;
- **foreach** *candidate* $t \in C_i$ **do**
  - **if** $\mathrm{wells}(t)$ *is contained in concatenation (forward or reverse) of any pair in* $\mathcal{O}$ **then**
    - Remove $t$ from $C_i$;
- Replenish $C_i$ from backup if needed;
- $\mathcal{T}[i] \leftarrow$ best remaining candidate in $C_i$;

`// Step 4: Resolve overlaps between consecutive positions`

**for** $i = 0$ **to** $|\mathcal{T}| - 2$ **do**
- **if** *last two wells of* $\mathcal{T}[i]$ *equal first two wells of* $\mathcal{T}[i+1]$ **then**
  - $p \leftarrow$ argmin-count among $\{i, i+1\}$;
  - Replace $\mathcal{T}[p]$ with best alternative in $C_p$ avoiding overlap;

**return** $\mathcal{T}$;

**Algorithm 7:** De Bruijn Graph Assembly for Multiplex Chain Recovery

```
Input: Fragment set 𝓕 (forward + reverse), k-mer size k, start anchors A_s, end anchors A_e, min edge count μ,
       beam width β, target length n
Output: Assembled oligo-ID sequences 𝓢
// Step 1: k-mer counting
Initialise token map φ : token ↔ ℤ;
𝒦 ← ∅;
foreach fragment f ∈ 𝓕 do
    x ← φ.encode(f);
    for i = 0 to |x| − k do
        𝒦[x[i..i+k]] ← 𝒦[x[i..i+k]] + 1;
// Step 2: Build de Bruijn graph
Initialise directed graph G = (V, E);
foreach k-mer κ ∈ 𝒦 do
    Add edge (κ[1 : k−1] → κ[2 : k]) with weight 𝒦[κ];
// Step 3: Prune weak edges
foreach edge (u, v) ∈ E do
    if w(u, v) < μ then remove (u, v) from G;
// Step 4: Anchor resolution
foreach anchor pair (a_s^(i), a_e^(i)) do
    V_s ← nodes matching suffix of φ(a_s^(i));
    V_e ← nodes matching suffix of φ(a_e^(i));
    if V_s = ∅ then V_s ← {v ∈ V : in(v) = 0};
    if V_e = ∅ then V_e ← {v ∈ V : out(v) = 0};
    // Step 5: Beam search with backtracking
    𝓑 ← {([s], seq(s), 0) : s ∈ V_s};
    𝓡 ← ∅;
    while 𝓑 ≠ ∅ do
        𝓑′ ← ∅;
        foreach path (P, x, c) ∈ 𝓑 do
            if |x| = n and x ends with φ(a_e^(i)) then
                𝓡 ← 𝓡 ∪ {(P, x, c)};
                continue;
            if x ends with φ(a_e^(i)) and |x| < n then
                continue;
            v ← last node in P;
            foreach (v, u) ∈ E sorted by weight (top β) do
                if u ∈ P then continue;
                𝓑′ ← 𝓑′ ∪ {(P · u, seq(u), c − log w(v, u))};
        Sort 𝓑′ by score; keep top β;
        𝓑 ← 𝓑′;
    Store 𝓡^(i) ← {φ^−1(x) : (P, x, c) ∈ 𝓡};
return {𝓡^(1), ..., 𝓡^(|𝒜|)};
```

#### Supplementary Note 6: Binary

This section presents the binary input data used for the encoding process.
CE:
0110101010110001111111110000010010010001000110110001000010101000000011111110000100010100010
0000010100000000101111100010000100010100100010001000000001000001000001000000010001111111110
1010101010101011111111110
CCE:
0000111001101101100111000000000101011011011010100000001000010111111111101000000111000010000 1
1000010000000001000000100100100100001000000000000010100101000011100000000000100110010000001000
0001000111110011111000000000111001100011000110000000000100001111001111000000000000000000000111
1110000000001000000000010000100100000110000000000100001011100000100000100001000010010001000
0000111001100001100001110010000100010000000100000100011000011111111111111000000100000111111 1
1111111000000000000000110000000011000000000000001000000000010000100000000010000000000010001110
0010001000000000000010001000111010000000000000001000000010011000000000000011000000000101111111
1111111010000000010010010110100100100000001101001001100100101100000110010010011001001001100
0111001001001100100100111010011001001011010010011001110011111111111111110011011000000000 00
000000000011000111111111111111111111100
PE:
10101011111111011011001000100110110001110001010100011100001110000101000010100011011001000 10
1111111101010110101010111110
CPE:
0011001010100110000011010101011000000011111111000001001000100010000010001010101001000000000
1101100111000000001010100010010000111011100000100011001001100000000011101110000000000001111 10
00001000000010001000000100000001000100000000001001000100010000010010001000100000000111111000000
00011100011100000000110000011000000001000000010010010010000000100100101100000001100000001100
0000011000001111111111111000010101010101010000101010101010100011010101010101100110101010101
0110101010101010101011001111111111100101000000000000001001111111111111110

### Supplementary Note 7: Assembly protocols

#### A. CE assembly (208 bit)

Ligation step 1

| 1.1 | 1.2 | 1.3 | 1.4 | 1.5 | 1.6 | 1.7 | 1.8 | 1.9 | 1.10 |
|---|---|---|---|---|---|---|---|---|---|
| D/e* | E/b | A*/f* | A*/d | A/a* | B/h | C/c | A*/h* | A/a* | F*/e |
| F/e | E*/a | A/h* | A/h | A*/b* | E*/h* | B*/c* | E*/h | B/a | F/h |
| F*/a | A/a* | F*/h | H*/h* | B/b | E/d* | B/a* | E/a* | B*/h | H/h* |
| H/a* | A*/f | F/d* | H/a | B*/h* | C*/d | A/a | | H/h* | H*/g |
| H*/h | | | | E/h | | A*/h* | | H*/e* | |
| A*/h* | | | | | | | | F*/e | |
| A/b* | | | | | | | | | |

Ligation step 2

| 2.1 | 2.2 | 2.3 | 2.4 |
|---|---|---|---|
| 1.1 | 1.5 | 1.7 | A*/a |
| 1.2 | E*/h* | A/h | 1.9 |
| 1.3 | B*/h | 1.8 | F/e* |
| 1.4 | 1.6 | | 1.10 |

Purification 1

Ligation step 3

| 3.1 |
|---|
| 2.1 |
| 2.2 |
| 2.3 |
| 2.4 |

Purification 2

#### B. CCE assembly (832 bits)

**Codeword 1**

Ligation step 1

| 1.1 | 1.2 | 1.3 | 1.4 | 1.5 | 1.6 | 1.7 | 1.8 | 1.9 | 1.10 |
|---|---|---|---|---|---|---|---|---|---|
| D/b* | D*/d | C/e* | G/g* | F*/f* | D*/h* | A*/f | C/a | A*/b | G/f* |
| H/b | D/e | C*/a | G*/a | C/f | A*/h | A/a | B*/a* | A/e* | E*/f |
| H*/d* | G/e* | G/a* | B/a* | C*/g* | A/a* | C*/a* | B/c* | H*/e | E/b* |
| | G*/f* | G*/b | B*/f* | D/g | C*/a | | A/c | H/f | B*/b |
| | A/f | F*/b* | F/f | | C/f* | | A*/g* | G*/f* | B/d* |
| | A*/b | F/g | | | | | G/g | | |
| | H*/b* | | | | | | G*/b* | | |
| | H/e | | | | | | | | |

Ligation step 2

| 2.1 | 2.2 | 2.3 |
|---|---|---|
| 1.1 | 1.4 | 1.8 |
| 1.2 | 1.5 | 1.9 |
| 1.3 | 1.6 | 1.10 |
| | 1.7 | |

Purification 1

Ligation step 3

| 3.1 |
|---|
| 2.1 |
| 2.2 |
| 2.3 |

Purification 2

---

**Codeword 2**

Ligation step 1

| 1.1 | 1.2 | 1.3 | 1.4 | 1.5 | 1.6 | 1.7 | 1.8 | 1.9 | 1.10 | 1.11 |
|---|---|---|---|---|---|---|---|---|---|---|
| G*/d* | F*/c | D/b | H*/e | A/a | G/g | H/h | A*/a* | A/g | C/a* | F/h* |

| | | | | | | | | | | | |
|---|---|---|---|---|---|---|---|---|---|---|---|
| | B/d | E*/c* | B*/b* | D*/e* | C*/a* | B*/g* | E*/h* | A/b | A*/d | C*/c* | F*/e* |
| | B*/c* | E/b* | B/e | D/a | C/h* | B/a* | E/e | D/b* | G*/d* | F/c | H/e |
| | F/c | D*/b | H/e* | A*/a* | G*/h | H*/a | C/e* | D*/c | G/h | F*/h | H*/f |
| | | | | | | | C*/f* | C/c* | C/h* | | |
| | | | | | | | A*/f | C*/g* | C*/a | | |
| | | | | | | | A/a | | | | |
| Ligation step 2 | | | | | | | | | | | |
| | **2.1** | **2.2** | **2.3** | | | | | | | | |
| | 1.1 | 1.5 | 1.9 | | | | | | | | |
| | 1.2 | 1.6 | 1.10 | | | | | | | | |
| | 1.3 | 1.7 | 1.11 | | | | | | | | |
| | 1.4 | 1.8 | | | | | | | | | |
| Purification 1 | | | | | | | | | | | |
| Ligation step 3 | | | | | | | | | | | |
| | **3.1** | | | | | | | | | | |
| | 2.1 | | | | | | | | | | |
| | 2.2 | | | | | | | | | | |
| | 2.3 | | | | | | | | | | |
| Purification 2 | | | | | | | | | | | |
| **Codeword 3** | | | | | | | | | | | |
| Ligation step 1 | | | | | | | | | | | |
| | **1.1** | **1.2** | **1.3** | **1.4** | **1.5** | **1.6** | **1.7** | **1.8** | **1.9** | **1.10** | **1.11** |
| | E*/f* | E*/g | E*/a* | A/a | H/a* | A/d | A/h* | H/g* | D*/e* | B/h* | F*/g* |
| | H/f | E/d | E/g* | E/a* | C*/a | A*/a* | A*/g* | D*/g | D/h* | A*/h | H/g |
| | H*/g* | A*/d* | H*/g | E*/h | C/f | E*/a | H/g | D/d* | B*/h | A/c* | H*/h* |
| | | A/a | H/f* | H*/h* | B*/f* | E/h | H*/d | A*/d | | F/c | A*/h |
| | | | G*/f | | B/d* | | H*/d* | A/a | | | A/b |
| | | | G/e* | | | | | E/a* | | | |
| | | | A*/e | | | | | E*/e | | | |
| Ligation step 2 | | | | | | | | | | | |
| | **2.1** | **2.2** | **2.3** | **2.4** | | | | | | | |
| | 1.1 | 1.4 | 1.7 | 1.9 | | | | | | | |
| | 1.2 | 1.5 | 1.8 | 1.10 | | | | | | | |
| | 1.3 | 1.6 | | 1.11 | | | | | | | |
| Purification 1 | | | | | | | | | | | |
| Ligation step 3 | | | | | | | | | | | |
| | **3.1** | | | | | | | | | | |
| | 2.1 | | | | | | | | | | |
| | 2.2 | | | | | | | | | | |
| | 2.3 | | | | | | | | | | |
| | 2.4 | | | | | | | | | | |
| Purification 2 | | | | | | | | | | | |
| **Codeword 5** | | | | | | | | | | | |
| Ligation step 1 | | | | | | | | | | | |
| | **1.1** | **1.2** | **1.3** | **1.4** | **1.5** | **1.6** | **1.7** | **1.8** | **1.9** | **1.10** | **1.11** |
| | A/b | D*/f | F/h* | B/a | A*/f | E*/b | H/b* | G/g | G*/a | F/c | F/h* |
| | E*/b* | A*/f* | H*/h | D/a* | A/a* | A*/b* | C/b | C/g* | G/h* | C*/c* | F*/d* |
| | E/d | A/a | H/a* | D*/d* | G*/a | A/a | C*/g | C*/c | A/h | C/h | H*/d |
| | G/d* | F*/a* | B*/a | H/d | G/b | H*/a* | G*/g* | B*/c* | A*/c* | | H/a |
| | G*/e | | | H*/g* | E/b* | | | B/a* | F*/c | | |
| | D/e* | | | F*/g | | | | | | | |
| | | | | F/f* | | | | | | | |
| Ligation step 2 | | | | | | | | | | | |
| | **2.1** | **2.2** | **2.3** | **2.4** | | | | | | | |
| | 1.1 | 1.4 | 1.6 | 1.10 | | | | | | | |
| | 1.2 | 1.5 | 1.7 | 1.11 | | | | | | | |
| | 1.3 | | 1.8 | | | | | | | | |
| | | | 1.9 | | | | | | | | |

Purification 1

Ligation step 3

| **3.1** |
|---|
| 2.1 |
| 2.2 |
| 2.3 |
| 2.4 |

Purification 2

---

**Codeword 4**

Ligation step 1

| **1.1** | **1.2** | **1.3** | **1.4** | **1.5** | **1.6** | **1.7** | **1.8** | **1.9** | **1.10** |
|---|---|---|---|---|---|---|---|---|---|
| E/g | G/f | G*/h* | D*/f | H/g | A/f* | A*/e* | C*/f | G*/c | A/d* |
| G*/g* | G*/d* | G/a* | D/g | F/g* | A*/h | A/a* | B/f* | H*/c* | H/d |
| G/f* | A/d | D*/a | C*/g* | F*/a* | H/h* | H/a | B*/c | H/d | H*/c* |
| | A*/h | D/d | C/h | A*/a | H*/e | H*/f* | G/c* | A*/d* | D*/c |
| | | H*/d* | A*/h* | A/b* | | C/f | | | D/h* |
| | | H/f* | A/a* | D/b | | | | | |
| | | | H*/a | D*/h | | | | | |
| | | | | G/h* | | | | | |
| | | | | G*/f | | | | | |

Ligation step 2

| **2.1** | **2.2** | **2.3** | **2.4** |
|---|---|---|---|
| 1.1 | 1.3 | 1.5 | 1.8 |
| 1.2 | 1.4 | 1.6 | 1.9 |
| | | 1.7 | 1.10 |

Purification 1

Ligation step 3

| **3.1** |
|---|
| 2.1 |
| 2.2 |
| 2.3 |
| 2.4 |

Purification 2

#### C. PE assembly (119 bits)

Ligation step 1

| **1.1** | **1.2** | **1.3** | **1.4** | **1.5** | **1.6** | **1.7** | **1.8** | **1.9** | **1.10** |
|---|---|---|---|---|---|---|---|---|---|
| A/c* | B/f* | B*/c | C/f | C*/e | D/f | D*/e | E/c | E*/f | F/d |
| H*/c | A/f | E*/c* | F/f* | E*/e* | A/f* | A/e* | D*/c* | D*/f* | E/d* |
| H/a | A*/a* | E/b | F*/b* | E/c | A*/c* | A*/d | D/d* | D/e | E*/e* |
| **1.11** | **1.12** | **1.13** | **1.14** | **1.15** | **1.16** | **1.17** | | | |
| F*/d | G/e | G*/f | H/d* | H*/b* | A/c* | A*/h | | | |
| E/d* | E*/e* | E/f* | A/d | A/b | H*/c | F/h* | | | |
| E*/f | E/f* | E*/g | A*/g* | A*/h | H/h* | F*/a | | | |

Ligation step 2

| **2.1** | **2.2** | **2.3** | **2.4** |
|---|---|---|---|
| 1.1 | 1.5 | 1.9 | 1.14 |
| 1.2 | 1.6 | 1.10 | 1.15 |
| 1.3 | 1.7 | 1.11 | 1.16 |
| 1.4 | 1.8 | 1.12 | 1.17 |
| | | 1.13 | |

Purification 1

Ligation step 3

| **3.1** |
|---|
| 2.1 |
| 2.2 |
| 2.3 |

2.4

**D. CPE assembly (544 bits)**

**Codeword 1**
Ligation step 1

| **1.1** | **1.2** | **1.3** | **1.4** | **1.5** | **1.6** | **1.7** | **1.8** | **1.9** | **1.10** |
|---|---|---|---|---|---|---|---|---|---|
| A/d | B/d | B*/a* | C/d | C*/a | D/a* | D*/f | E/a* | E*/d | F/a* |
| G/d* | D*/d* | G*/a | B/d* | H/a* | E/a | G*/f* | D*/a | B*/d* | C/a |
| G*/a | D/a* | G/b | B*/b* | H*/c | E*/c* | G/d | D/d* | B/e | C*/e* |
| **1.11** | **1.12** | **1.13** | **1.14** | **1.15** | **1.16** | **1.17** | | | |
| F*/d | G/c | G*/a* | H/f | H*/d | A/c | A*/d | | | |
| H/d* | A*/c* | H*/a | C/f* | D*/d* | F/c* | D*/d* | | | |
| H*/f | A/f* | H/g | C*/g* | D/h | F*/h* | D/a | | | |

**Codeword 2**
Ligation step 1

| **1.1** | **1.2** | **1.3** | **1.4** | **1.5** | **1.6** | **1.7** | **1.8** | **1.9** | **1.10** |
|---|---|---|---|---|---|---|---|---|---|
| A/b* | B/f | B*/d | C/c | C*/h | D/f | D*/c | E/a* | E*/d | F/a* |
| D*/b | H*/f* | C*/d* | G/c* | E*/h* | H*/f* | C*/c* | H/a | C*/d* | C*/a |
| D/a | H/a* | C/b | G*/b* | E/c | H/c* | C/d | H*/d* | C/e | C/e* |
| **1.11** | **1.12** | **1.13** | **1.14** | **1.15** | **1.16** | **1.17** | | | |
| F*/c | G/e | G*/d | H/a | H*/d | A/g | A*/d | | | |
| D/c* | A*/e* | F/d* | E*/a* | C*/d* | B*/g* | H*/d* | | | |
| D*/f | A/f* | F*/g | E/g* | C/h | B/h* | H/a | | | |

**Codeword 3**
Ligation step 1

| **1.1** | **1.2** | **1.3** | **1.4** | **1.5** | **1.6** | **1.7** | **1.8** | **1.9** | **1.10** |
|---|---|---|---|---|---|---|---|---|---|
| A/c* | B/b* | B*/d* | C/e | C*/b* | D/b* | D*/h | E/c | E*/c | F/d* |
| F/c | H*/b | E*/d | H/e* | E/b | H/b | C*/h* | B*/c* | F*/c* | C*/d |
| F*/a | H/a* | E/b | H*/b* | E*/c | H*/c* | C/d | B/d* | F/e | C/e* |
| **1.11** | **1.12** | **1.13** | **1.14** | **1.15** | **1.16** | **1.17** | | | |
| F*/e | G/d* | G*/c | H/h | H*/g* | A/g | A*/f | | | |
| C*/e* | F*/d | B/c* | A/h* | D/g | E*/g* | G/f* | | | |
| C/f | F/f* | B*/g | A*/g* | D*/h | E/h* | G*/a | | | |

**Codeword 4**
Ligation step 1

| **1.1** | **1.2** | **1.3** | **1.4** | **1.5** | **1.6** | **1.7** | **1.8** | **1.9** | **1.10** |
|---|---|---|---|---|---|---|---|---|---|
| A/f* | B/c* | B*/e* | C/c* | C*/g* | D/d* | D*/f* | E/b* | E*/b* | F/h* |
| H/f | C*/c | E*/e | G*/c | A*/g | E*/d | F*/f | F*/b | A*/b | A/h |
| H*/a | C/a* | E/b | G/b* | A/c | E/c* | F/d | F/d* | A/e | A*/e* |
| **1.11** | **1.12** | **1.13** | **1.14** | **1.15** | **1.16** | **1.17** | | | |
| F*/b* | G/g | G*/e | H/h | H*/b | A/g | A*/b* | | | |
| A*/b | C/g* | H*/e* | C*/h* | A/b* | F/g* | B/b | | | |
| A/f | C*/f* | H/g | C/g* | A*/h | F*/h* | B*/a | | | |

**Codeword 5**
Ligation step 1

| **1.1** | **1.2** | **1.3** | **1.4** | **1.5** | **1.6** | **1.7** | **1.8** | **1.9** | **1.10** |
|---|---|---|---|---|---|---|---|---|---|
| A/g* | B/g* | B*/g* | C/c* | C*/h* | D/d* | D*/h* | E/g | E*/g* | F/b |
| D/g | C*/g | A*/g | B*/c | A/h | F*/d | A*/h | F*/g* | H*/g | C/b* |
| D*/a | C/a* | A/b | B/b* | A*/c | F/c* | A/d | F/d* | H/e | C*/e* |
| **1.11** | **1.12** | **1.13** | **1.14** | **1.15** | **1.16** | **1.17** | | | |
| F*/e* | G/b | G*/b | H/h* | H*/b | A/b | A*/g | | | |
| E*/e | D/b* | A/b* | G/h | C/b* | H/b* | C/g* | | | |
| E/f | D*/f* | A*/g | G*/g* | C*/h | H*/h* | C*/a | | | |

**All codewords in the same pool**
Ligation step 2

| **2.1** | **2.2** | **2.3** | **2.4** |
|---|---|---|---|

| | | | | |
|---|---|---|---|---|
| | 1.1 | 1.6 | 1.10 | 1.14 |
| | 1.2 | 1.7 | 1.11 | 1.15 |
| | 1.3 | 1.8 | 1.12 | 1.16 |
| | 1.4 | 1.9 | 1.13 | 1.17 |
| | 1.5 | | | |
| Purification 1 | | | | |
| Ligation step 3 | | | | |
| | **3.1** | | | |
| | 2.1 | | | |
| | 2.2 | | | |
| | 2.3 | | | |
| | 2.4 | | | |
| Purification 2 | | | | |

#### Supplementary Note 8: DNA Strands

In this section, we list all ordered oligonucleotides. Asterisks on motif labels indicate reverse complements.

##### A. Library strands

**Table 5.** Library oligonucleotoide strands and their domain motif structure

| Index Number | Motif composition | Sequence |
|---|---|---|
| 1 | A/a | TGCTAAGACGACCGGAAAAG |
| 2 | A/a* | TGCTAAGACGCTTTTCCGGT |
| 3 | A/b | TGCTAAGACGCCGGAGTTTT |
| 4 | A/b* | TGCTAAGACGAAAACTCCGG |
| 5 | A/c | TGCTAAGACGCATTGCGAAC |
| 6 | A/c* | TGCTAAGACGGTTCGCAATG |
| 7 | A/d | TGCTAAGACGCGCAACTAGA |
| 8 | A/d* | TGCTAAGACGTCTAGTTGCG |
| 9 | A/e | TGCTAAGACGGCATAACCGT |
| 10 | A/e* | TGCTAAGACGACGGTTATGC |
| 11 | A/f | TGCTAAGACGCGAAGTACGA |
| 12 | A/f* | TGCTAAGACGTCGTACTTCG |
| 13 | A/g | TGCTAAGACGCGTTGAATGC |
| 14 | A/g* | TGCTAAGACGGCATTCAACG |
| 15 | A/h | TGCTAAGACGATGGAATCGC |
| 16 | A/h* | TGCTAAGACGGCGATTCCAT |
| 17 | A*/a | CGTCTTAGCAACCGGAAAAG |
| 18 | A*/a* | CGTCTTAGCACTTTTCCGGT |
| 19 | A*/b | CGTCTTAGCACCGGAGTTTT |
| 20 | A*/b* | CGTCTTAGCAAAAACTCCGG |
| 21 | A*/c | CGTCTTAGCACATTGCGAAC |
| 22 | A*/c* | CGTCTTAGCAGTTCGCAATG |
| 23 | A*/d | CGTCTTAGCACGCAACTAGA |
| 24 | A*/d* | CGTCTTAGCATCTAGTTGCG |
| 25 | A*/e | CGTCTTAGCAGCATAACCGT |
| 26 | A*/e* | CGTCTTAGCAACGGTTATGC |
| 27 | A*/f | CGTCTTAGCACGAAGTACGA |
| 28 | A*/f* | CGTCTTAGCATCGTACTTCG |
| 29 | A*/g | CGTCTTAGCACGTTGAATGC |
| 30 | A*/g* | CGTCTTAGCAGCATTCAACG |
| 31 | A*/h | CGTCTTAGCAATGGAATCGC |
| 32 | A*/h* | CGTCTTAGCAGCGATTCCAT |
| 33 | B/a | TAGCGATCCTACCGGAAAAG |
| 34 | B/a* | TAGCGATCCTCTTTTCCGGT |
| 35 | B/b | TAGCGATCCTCCGGAGTTTT |
| 36 | B/b* | TAGCGATCCTAAAACTCCGG |
| 37 | B/c | TAGCGATCCTCATTGCGAAC |
| 38 | B/c* | TAGCGATCCTGTTCGCAATG |
| 39 | B/d | TAGCGATCCTCGCAACTAGA |
| 40 | B/d* | TAGCGATCCTTCTAGTTGCG |
| 41 | B/e | TAGCGATCCTGCATAACCGT |
| 42 | B/e* | TAGCGATCCTACGGTTATGC |
| 43 | B/f | TAGCGATCCTCGAAGTACGA |
| 44 | B/f* | TAGCGATCCTTCGTACTTCG |
| 45 | B/g | TAGCGATCCTCGTTGAATGC |
| 46 | B/g* | TAGCGATCCTGCATTCAACG |
| 47 | B/h | TAGCGATCCTATGGAATCGC |
| 48 | B/h* | TAGCGATCCTGCGATTCCAT |
| 49 | B*/a | AGGATCGCTAACCGGAAAAG |
| 50 | B*/a* | AGGATCGCTACTTTTCCGGT |

| | | |
|---|---|---|
| 51 | B*/b | AGGATCGCTACCGGAGTTTT |
| 52 | B*/b* | AGGATCGCTAAAAACTCCGG |
| 53 | B*/c | AGGATCGCTACATTGCGAAC |
| 54 | B*/c* | AGGATCGCTAGTTCGCAATG |
| 55 | B*/d | AGGATCGCTACGCAACTAGA |
| 56 | B*/d* | AGGATCGCTATCTAGTTGCG |
| 57 | B*/e | AGGATCGCTAGCATAACCGT |
| 58 | B*/e* | AGGATCGCTAACGGTTATGC |
| 59 | B*/f | AGGATCGCTACGAAGTACGA |
| 60 | B*/f* | AGGATCGCTATCGTACTTCG |
| 61 | B*/g | AGGATCGCTACGTTGAATGC |
| 62 | B*/g* | AGGATCGCTAGCATTCAACG |
| 63 | B*/h | AGGATCGCTAATGGAATCGC |
| 64 | B*/h* | AGGATCGCTAGCGATTCCAT |
| 65 | C/a | CTCACGCTAAACCGGAAAAG |
| 66 | C/a* | CTCACGCTAACTTTTCCGGT |
| 67 | C/b | CTCACGCTAACCGGAGTTTT |
| 68 | C/b* | CTCACGCTAAAAAACTCCGG |
| 69 | C/c | CTCACGCTAACATTGCGAAC |
| 70 | C/c* | CTCACGCTAAGTTCGCAATG |
| 71 | C/d | CTCACGCTAACGCAACTAGA |
| 72 | C/d* | CTCACGCTAATCTAGTTGCG |
| 73 | C/e | CTCACGCTAAGCATAACCGT |
| 74 | C/e* | CTCACGCTAAACGGTTATGC |
| 75 | C/f | CTCACGCTAACGAAGTACGA |
| 76 | C/f* | CTCACGCTAATCGTACTTCG |
| 77 | C/g | CTCACGCTAACGTTGAATGC |
| 78 | C/g* | CTCACGCTAAGCATTCAACG |
| 79 | C/h | CTCACGCTAAATGGAATCGC |
| 80 | C/h* | CTCACGCTAAGCGATTCCAT |
| 81 | C*/a | TTAGCGTGAGACCGGAAAAG |
| 82 | C*/a* | TTAGCGTGAGCTTTTCCGGT |
| 83 | C*/b | TTAGCGTGAGCCGGAGTTTT |
| 84 | C*/b* | TTAGCGTGAGAAAACTCCGG |
| 85 | C*/c | TTAGCGTGAGCATTGCGAAC |
| 86 | C*/c* | TTAGCGTGAGGTTCGCAATG |
| 87 | C*/d | TTAGCGTGAGCGCAACTAGA |
| 88 | C*/d* | TTAGCGTGAGTCTAGTTGCG |
| 89 | C*/e | TTAGCGTGAGGCATAACCGT |
| 90 | C*/e* | TTAGCGTGAGACGGTTATGC |
| 91 | C*/f | TTAGCGTGAGCGAAGTACGA |
| 92 | C*/f* | TTAGCGTGAGTCGTACTTCG |
| 93 | C*/g | TTAGCGTGAGCGTTGAATGC |
| 94 | C*/g* | TTAGCGTGAGGCATTCAACG |
| 95 | C*/h | TTAGCGTGAGATGGAATCGC |
| 96 | C*/h* | TTAGCGTGAGGCGATTCCAT |
| 97 | D/a | AAAGAACCCGACCGGAAAAG |
| 98 | D/a* | AAAGAACCCGCTTTTCCGGT |
| 99 | D/b | AAAGAACCCGCCGGAGTTTT |
| 100 | D/b* | AAAGAACCCGAAAACTCCGG |
| 101 | D/c | AAAGAACCCGCATTGCGAAC |
| 102 | D/c* | AAAGAACCCGGTTCGCAATG |
| 103 | D/d | AAAGAACCCGCGCAACTAGA |
| 104 | D/d* | AAAGAACCCGTCTAGTTGCG |
| 105 | D/e | AAAGAACCCGGCATAACCGT |
| 106 | D/e* | AAAGAACCCGACGGTTATGC |
| 107 | D/f | AAAGAACCCGCGAAGTACGA |
| 108 | D/f* | AAAGAACCCGTCGTACTTCG |

| | | |
|---|---|---|
| 109 | D/g | AAAGAACCCGCGTTGAATGC |
| 110 | D/g* | AAAGAACCCGGCATTCAACG |
| 111 | D/h | AAAGAACCCGATGGAATCGC |
| 112 | D/h* | AAAGAACCCGGCGATTCCAT |
| 113 | D*/a | CGGGTTCTTTACCGGAAAAG |
| 114 | D*/a* | CGGGTTCTTTCTTTTCCGGT |
| 115 | D*/b | CGGGTTCTTTCCGGAGTTTT |
| 116 | D*/b* | CGGGTTCTTTAAAACTCCGG |
| 117 | D*/c | CGGGTTCTTTCATTGCGAAC |
| 118 | D*/c* | CGGGTTCTTTGTTCGCAATG |
| 119 | D*/d | CGGGTTCTTTCGCAACTAGA |
| 120 | D*/d* | CGGGTTCTTTTCTAGTTGCG |
| 121 | D*/e | CGGGTTCTTTGCATAACCGT |
| 122 | D*/e* | CGGGTTCTTTACGGTTATGC |
| 123 | D*/f | CGGGTTCTTTCGAAGTACGA |
| 124 | D*/f* | CGGGTTCTTTTCGTACTTCG |
| 125 | D*/g | CGGGTTCTTTCGTTGAATGC |
| 126 | D*/g* | CGGGTTCTTTGCATTCAACG |
| 127 | D*/h | CGGGTTCTTTATGGAATCGC |
| 128 | D*/h* | CGGGTTCTTTGCGATTCCAT |
| 129 | E/a | GCGTACCATTACCGGAAAAG |
| 130 | E/a* | GCGTACCATTCTTTTCCGGT |
| 131 | E/b | GCGTACCATTCCGGAGTTTT |
| 132 | E/b* | GCGTACCATTAAAACTCCGG |
| 133 | E/c | GCGTACCATTCATTGCGAAC |
| 134 | E/c* | GCGTACCATTGTTCGCAATG |
| 135 | E/d | GCGTACCATTCGCAACTAGA |
| 136 | E/d* | GCGTACCATTTCTAGTTGCG |
| 137 | E/e | GCGTACCATTGCATAACCGT |
| 138 | E/e* | GCGTACCATTACGGTTATGC |
| 139 | E/f | GCGTACCATTCGAAGTACGA |
| 140 | E/f* | GCGTACCATTTCGTACTTCG |
| 141 | E/g | GCGTACCATTCGTTGAATGC |
| 142 | E/g* | GCGTACCATTGCATTCAACG |
| 143 | E/h | GCGTACCATTATGGAATCGC |
| 144 | E/h* | GCGTACCATTGCGATTCCAT |
| 145 | E*/a | AATGGTACGCACCGGAAAAG |
| 146 | E*/a* | AATGGTACGCCTTTTCCGGT |
| 147 | E*/b | AATGGTACGCCCGGAGTTTT |
| 148 | E*/b* | AATGGTACGCAAAACTCCGG |
| 149 | E*/c | AATGGTACGCCATTGCGAAC |
| 150 | E*/c* | AATGGTACGCGTTCGCAATG |
| 151 | E*/d | AATGGTACGCCGCAACTAGA |
| 152 | E*/d* | AATGGTACGCTCTAGTTGCG |
| 153 | E*/e | AATGGTACGCGCATAACCGT |
| 154 | E*/e* | AATGGTACGCACGGTTATGC |
| 155 | E*/f | AATGGTACGCCGAAGTACGA |
| 156 | E*/f* | AATGGTACGCTCGTACTTCG |
| 157 | E*/g | AATGGTACGCCGTTGAATGC |
| 158 | E*/g* | AATGGTACGCGCATTCAACG |
| 159 | E*/h | AATGGTACGCATGGAATCGC |
| 160 | E*/h* | AATGGTACGCGCGATTCCAT |
| 161 | F/a | GATGCGGAATACCGGAAAAG |
| 162 | F/a* | GATGCGGAATCTTTTCCGGT |
| 163 | F/b | GATGCGGAATCCGGAGTTTT |
| 164 | F/b* | GATGCGGAATAAAACTCCGG |
| 165 | F/c | GATGCGGAATCATTGCGAAC |
| 166 | F/c* | GATGCGGAATGTTCGCAATG |

| | | |
|---|---|---|
| 167 | F/d | GATGCGGAATCGCAACTAGA |
| 168 | F/d* | GATGCGGAATTCTAGTTGCG |
| 169 | F/e | GATGCGGAATGCATAACCGT |
| 170 | F/e* | GATGCGGAATACGGTTATGC |
| 171 | F/f | GATGCGGAATCGAAGTACGA |
| 172 | F/f* | GATGCGGAATTCGTACTTCG |
| 173 | F/g | GATGCGGAATCGTTGAATGC |
| 174 | F/g* | GATGCGGAATGCATTCAACG |
| 175 | F/h | GATGCGGAATATGGAATCGC |
| 176 | F/h* | GATGCGGAATGCGATTCCAT |
| 177 | F*/a | ATTCCGCATCACCGGAAAAG |
| 178 | F*/a* | ATTCCGCATCCTTTTCCGGT |
| 179 | F*/b | ATTCCGCATCCCGGAGTTTT |
| 180 | F*/b* | ATTCCGCATCAAAACTCCGG |
| 181 | F*/c | ATTCCGCATCCATTGCGAAC |
| 182 | F*/c* | ATTCCGCATCGTTCGCAATG |
| 183 | F*/d | ATTCCGCATCCGCAACTAGA |
| 184 | F*/d* | ATTCCGCATCTCTAGTTGCG |
| 185 | F*/e | ATTCCGCATCGCATAACCGT |
| 186 | F*/e* | ATTCCGCATCACGGTTATGC |
| 187 | F*/f | ATTCCGCATCCGAAGTACGA |
| 188 | F*/f* | ATTCCGCATCTCGTACTTCG |
| 189 | F*/g | ATTCCGCATCCGTTGAATGC |
| 190 | F*/g* | ATTCCGCATCGCATTCAACG |
| 191 | F*/h | ATTCCGCATCATGGAATCGC |
| 192 | F*/h* | ATTCCGCATCGCGATTCCAT |
| 193 | G/a | TATCCAGGCAACCGGAAAAG |
| 194 | G/a* | TATCCAGGCACTTTTCCGGT |
| 195 | G/b | TATCCAGGCACCGGAGTTTT |
| 196 | G/b* | TATCCAGGCAAAAACTCCGG |
| 197 | G/c | TATCCAGGCACATTGCGAAC |
| 198 | G/c* | TATCCAGGCAGTTCGCAATG |
| 199 | G/d | TATCCAGGCACGCAACTAGA |
| 200 | G/d* | TATCCAGGCATCTAGTTGCG |
| 201 | G/e | TATCCAGGCAGCATAACCGT |
| 202 | G/e* | TATCCAGGCAACGGTTATGC |
| 203 | G/f | TATCCAGGCACGAAGTACGA |
| 204 | G/f* | TATCCAGGCATCGTACTTCG |
| 205 | G/g | TATCCAGGCACGTTGAATGC |
| 206 | G/g* | TATCCAGGCAGCATTCAACG |
| 207 | G/h | TATCCAGGCAATGGAATCGC |
| 208 | G/h* | TATCCAGGCAGCGATTCCAT |
| 209 | G*/a | TGCCTGGATAACCGGAAAAG |
| 210 | G*/a* | TGCCTGGATACTTTTCCGGT |
| 211 | G*/b | TGCCTGGATACCGGAGTTTT |
| 212 | G*/b* | TGCCTGGATAAAAACTCCGG |
| 213 | G*/c | TGCCTGGATACATTGCGAAC |
| 214 | G*/c* | TGCCTGGATAGTTCGCAATG |
| 215 | G*/d | TGCCTGGATACGCAACTAGA |
| 216 | G*/d* | TGCCTGGATATCTAGTTGCG |
| 217 | G*/e | TGCCTGGATAGCATAACCGT |
| 218 | G*/e* | TGCCTGGATAACGGTTATGC |
| 219 | G*/f | TGCCTGGATACGAAGTACGA |
| 220 | G*/f* | TGCCTGGATATCGTACTTCG |
| 221 | G*/g | TGCCTGGATACGTTGAATGC |
| 222 | G*/g* | TGCCTGGATAGCATTCAACG |
| 223 | G*/h | TGCCTGGATAATGGAATCGC |
| 224 | G*/h* | TGCCTGGATAGCGATTCCAT |

| | | |
|---|---|---|
| 225 | H/a | TAAGTCAGCGACCGGAAAAG |
| 226 | H/a* | TAAGTCAGCGCTTTTCCGGT |
| 227 | H/b | TAAGTCAGCGCCGGAGTTTT |
| 228 | H/b* | TAAGTCAGCGAAAACTCCGG |
| 229 | H/c | TAAGTCAGCGCATTGCGAAC |
| 230 | H/c* | TAAGTCAGCGGTTCGCAATG |
| 231 | H/d | TAAGTCAGCGCGCAACTAGA |
| 232 | H/d* | TAAGTCAGCGTCTAGTTGCG |
| 233 | H/e | TAAGTCAGCGGCATAACCGT |
| 234 | H/e* | TAAGTCAGCGACGGTTATGC |
| 235 | H/f | TAAGTCAGCGCGAAGTACGA |
| 236 | H/f* | TAAGTCAGCGTCGTACTTCG |
| 237 | H/g | TAAGTCAGCGCGTTGAATGC |
| 238 | H/g* | TAAGTCAGCGGCATTCAACG |
| 239 | H/h | TAAGTCAGCGATGGAATCGC |
| 240 | H/h* | TAAGTCAGCGGCGATTCCAT |
| 241 | H*/a | CGCTGACTTAACCGGAAAAG |
| 242 | H*/a* | CGCTGACTTACTTTTCCGGT |
| 243 | H*/b | CGCTGACTTACCGGAGTTTT |
| 244 | H*/b* | CGCTGACTTAAAAACTCCGG |
| 245 | H*/c | CGCTGACTTACATTGCGAAC |
| 246 | H*/c* | CGCTGACTTAGTTCGCAATG |
| 247 | H*/d | CGCTGACTTACGCAACTAGA |
| 248 | H*/d* | CGCTGACTTATCTAGTTGCG |
| 249 | H*/e | CGCTGACTTAGCATAACCGT |
| 250 | H*/e* | CGCTGACTTAACGGTTATGC |
| 251 | H*/f | CGCTGACTTACGAAGTACGA |
| 252 | H*/f* | CGCTGACTTATCGTACTTCG |
| 253 | H*/g | CGCTGACTTACGTTGAATGC |
| 254 | H*/g* | CGCTGACTTAGCATTCAACG |
| 255 | H*/h | CGCTGACTTAATGGAATCGC |
| 256 | H*/h* | CGCTGACTTAGCGATTCCAT |

#### B. Gate strands

**Table 6.** Computational oligonucleotide strands and their domain motif structure

| Name | Motif composition<br>Sequence |
|---|---|
| AND Loop | D/f/G/g/E/d<br>AAAGAACCCGCGAAGTACGATATCCAGGCACGTTGAATGCGCGTACCATTCGCAACTAGA |
| AND Hold | E*/f*/D*/b*<br>AATGGTACGCTCGTACTTCGCGGGTTCTTTAAAACTCCGG |
| AND Tail | H/b<br>TAAGTCAGCGCCGGAGTTTT |
| AND Input1 | F*/b*/H*/c<br>ATTCCGCATCAAAACTCCGGCGCTGACTTACATTGCGAAC |
| AND Input 2 | F/b/D/f/E/d<br>GATGCGGAATCCGGAGTTTTAAAGAACCCGCGAAGTACGAGCGTACCATTCGCAACTAG<br>A |
| OR Loop | A/a/G/g/E/d<br>TGCTAAGACGACCGGAAAAGTATCCAGGCACGTTGAATGCGCGTACCATTCGCAACTAGA |
| OR Hold | H*/d*/E*/a*/A*/b<br>CGCTGACTTATCTAGTTGCGAATGGTACGCCTTTTCCGGTCGTCTTAGCAAAAACTCCGG |
| OR Input 1 | A/a/E/d/H/f<br>TGCTAAGACGACCGGAAAAGGCGTACCATTCGCAACTAGATAAGTCAGCGCGAAGTACG<br>A |
| OR Input 2 | D/b/A/a/E/d |

| | |
|---|---|
| | AAAGAACCCGCCGGAGTTTTTGCTAAGACGACCGGAAAAGGCGTACCATTCGCAACTAG A |
| Reporter flurecence | d*/E*/g*/G* TCTAGTTGCGAATGGTACGCGCATTCAACGTGCCTGGATA/FAM/ |
| Reporter quencer | G/g /BHQ_1/TATCCAGGCACGTTGAATGC |

#### Supplementary Note 9: References

1. Saul B. Needleman and Christian D. Wunsch. A general method applicable to the search for similarities in the amino acid sequence of two proteins. *Journal of Molecular Biology*, 48(3):443–453, 1970. ISSN 0022-2836. doi: https://doi.org/10.1016/0022-2836(70)90057-4. URL https://www.sciencedirect.com/science/article/pii/0022283670900574.
